# Structural and mechanistic insights into human choline and ethanolamine transport

**DOI:** 10.1101/2023.09.15.557925

**Authors:** Keiken Ri, Tsai-Hsuan Weng, Ainara Claveras Cabezudo, Wiebke Jösting, Zhang Yu, Andre Bazzone, Nancy C.P. Leong, Sonja Welsch, Raymond T. Doty, Gonca Gursu, Tiffany Jia Ying Lim, Sarah Luise Schmidt, Janis L. Abkowitz, Gerhard Hummer, Di Wu, Long N Nguyen, Schara Safarian

**Author notes:** Correspondence and requests for materials should be addressed to Gerhard Hummer, Di Wu, Long Nguyen, and Schara Safarian. These authors contributed equally (Keiken Ri, Tsai-Hsuan Weng, Ainara Claveras Cabezudo).

## Abstract

Human feline leukemia virus subgroup C receptor-related proteins 1 and 2 (FLVCR1 and 2) are members of the major facilitator superfamily^1^. Their dysfunction is linked to several clinical disorders, including PCARP, HSAN, and Fowler syndrome^2–7^. Earlier studies concluded that FLVCR1 may function as a putative heme exporter^8–12^, while FLVCR2 was suggested to act as a heme importer^13^, yet conclusive biochemical and detailed molecular evidence remained elusive for the function of both transporters^14–17^. Here, we show that FLVCR1 and FLVCR2 facilitate the transport of choline and ethanolamine across human plasma membranes, utilizing a concentration-driven substrate translocation process. Through structural and computational analyses, we have identified distinct conformational states of FLVCRs and unraveled the coordination chemistry underlying their substrate interactions. Within the binding pocket of both transporters, we identify fully conserved tryptophan and tyrosine residues holding a central role in the formation of cation-π interactions, essential for choline and ethanolamine selectivity. Our findings not only clarify the mechanisms of choline and ethanolamine transport by FLVCR1 and FLVCR2, enhancing our comprehension of disease-associated mutations that interfere with these vital processes, but also shed light on the conformational dynamics of these MFS-type proteins during the transport cycle.

## Introduction

The feline leukaemia virus subgroup C receptor (FLVCR) family, a member of the major facilitator superfamily (MFS) of secondary active transporters, consists of four paralogues encoded by the human *SLC49* gene group^1^. FLVCR1 (SLC49A1) was initially identified as the cell surface receptor for feline leukaemia virus (FeLV)^18^. FLVCR2 (SLC49A2) shares 60% sequence identity with FLVCR1 in the transmembrane domain but does not bind the feline leukaemia virus subgroup C envelope protein^19^. Both transporters exhibit ubiquitous tissue distribution in humans and have significant haemato- and neuropathological implications^1,17^. Dysfunction of FLVCR1 caused by germline mutations is associated with posterior column ataxia with retinitis pigmentosa (PCARP)^2,3^, and hereditary sensory and autonomic neuropathies (HSANs)^4,5^. Similarly, truncation and missense mutations in *FLVCR2* are associated with autosomal-recessive cerebral proliferative vasculopathy (Fowler syndrome)^6,7^. Furthermore, both FLVCR variants are suggested to play a key role in cell development and differentiation, including angiogenesis and tumorigenesis^20–23^.

Earlier studies concluded that FLVCR1 may function as a putative heme exporter^8–12^, while FLVCR2 was suggested to act as a heme importer^13^, yet their definitive roles in this capacity remain elusive^14–17^. To accurately understand their functions, experimental validation at the biochemical and molecular levels is necessary, which will connect these transporters’ physiological roles and clinical relevance to their specific mechanistic actions. Recent studies indicated FLVCR1 is involved in choline transport, however, the ligands for FLVCR2 still remain elusive^24^. Here, we used an integrative approach, including cell-based radioligand transport assays, single-particle analysis (SPA) cryogenic-electron microscopy (cryo-EM), structure-guided mutagenesis, and atomistic molecular dynamics (MD) simulations to characterize the ligand specificity, molecular architecture, and the conformational landscape of FLVCR1 and FLVCR2 transporters.

## Results

### Choline and ethanolamine are transport substates of FLVCR1 and FLVCR2

We expressed the human *FLVCR1* and *FLVCR2* genes in human embryonic kidney (HEK293) cells to substantiate and characterize their roles in cellular choline and ethanolamine transport^25,26^. Upon overproduction, FLVCR1a (hereafter referred to as FLVCR1) and FLVCR2 localized at the plasma membrane of HEK293 cells (Fig. 1a and Supplementary Fig. 1a-d). Radioactive [^3^H] choline transport assays showed a discernible increase in uptake facilitated by the FLVCR2, whereas FLVCR1 did not exhibit such an effect under the tested condition (Extended Data Fig. 1a). Notably, co-expression of the choline kinase A (*CHKA*) gene significantly enhanced choline uptake by both transporters in dose- and time-dependent manners (Fig. 1b-d). This suggests that choline influx mediated by FLVCR1 and FLVCR2 is enhanced by the overproduction of downstream choline-utilizing enzymes. In order to ascertain whether choline is the principal physiological substrate for FLVCR1, a comprehensive metabolomic analysis was performed on liver samples from FLVCR1 knockout mice (Extended Data Fig. 2a). This led us to discover that, in addition to choline and its metabolites, ethanolamine metabolite profiles were also affected in FLVCR1 knockout livers (Extended Data Fig. 2b,c). Subsequent cell-based assays revealed that both FLVCRs facilitate ethanolamine uptake into cells (Extended Data Fig. 1b). Notably, co-expression of *ETNK-1* enhanced the ethanolamine transport rate of FLVCR1 by five-fold, but did not substantially affect the efficiency of FLVCR2 (Fig. 1e-g and Extended Data Fig. 1b). We determined transport kinetic parameters of FLVCR1 and FLVCR2, revealing *K_m_* values of 47.4 ± 9.8 µM and 64.0 ± 21.0 µM for choline, 8 ± 1.5 µM and 41.5 ± 32.0 µM for ethanolamine, respectively (Fig. 1c,f). To decipher the transport mechanism of both FLVCRs, we investigated their reliance on sodium ions and pH levels. Our uptake studies revealed that FLVCR-mediated transport of choline and ethanolamine is not contingent upon sodium ion involvement and operates effectively across a broad pH range (Extended Data Fig. 3a,b). This finding underscores that neither sodium nor pH gradients are critical for the translocation of choline and ethanolamine by FLVCRs. Next, we further assessed the mechanistic properties of FLVCR2 by performing a choline washout assay. Upon inverting the choline gradient across the plasma membrane, we measured a significant decrease of cellular choline levels within 1 h indicating for a bidirectional choline transport activity mediated by FLVCR2 (Extended Data Fig. 3c). FLVCR1 showed similar properties in an ethanolamine washout experiment (Extended Data Fig. 3c). Our findings suggest that both FLVCR1 and FLVCR2 operate as uniporters, facilitating downhill ligand transport independent of sodium or pH gradients but aided by the typically negative membrane potential^27^.

**Fig. 1:**
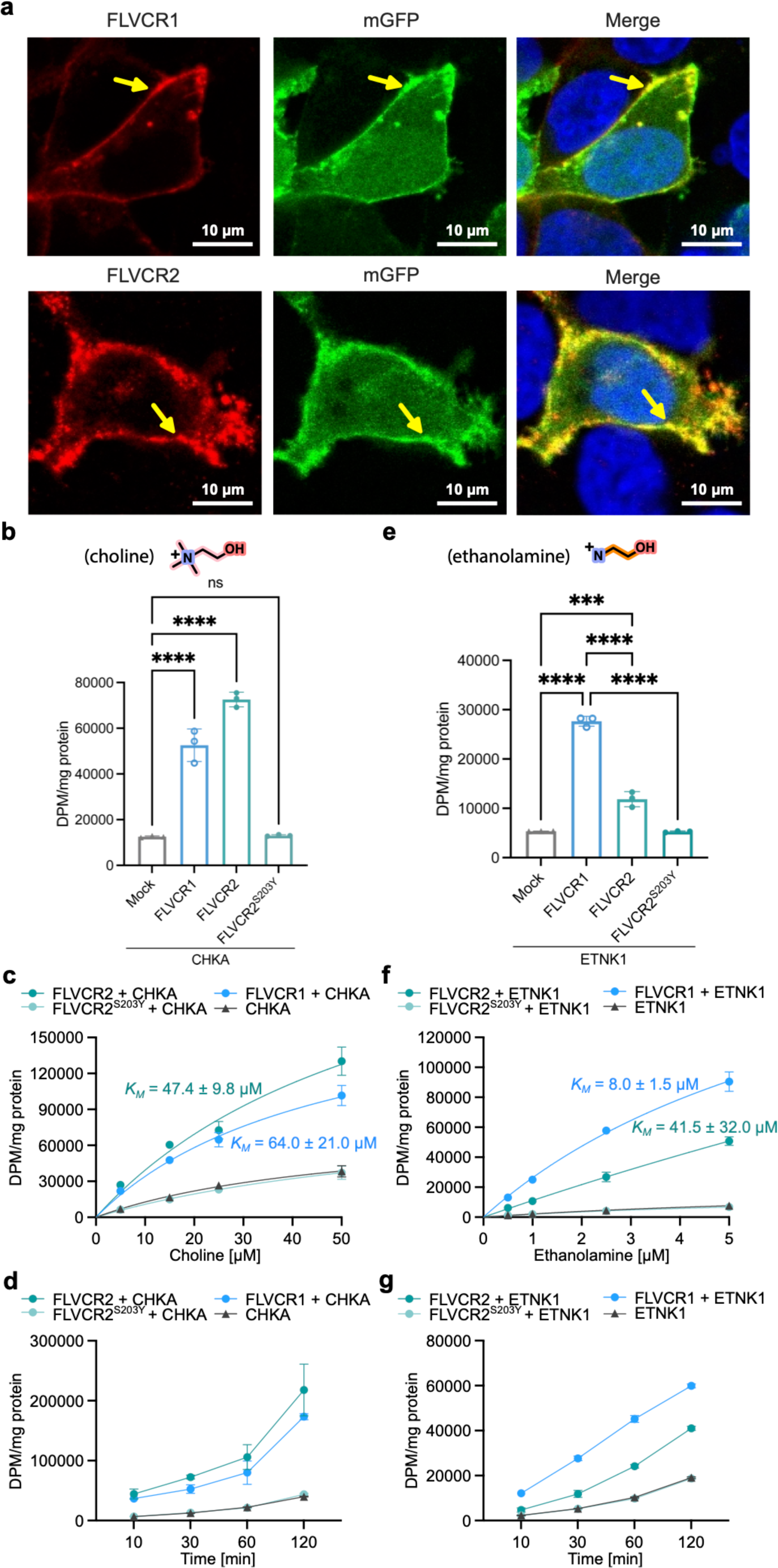
FLVCR1 and FLVCR2 are choline and ethanolamine transporters. **a**, Confocal imaging shows that FLVCR1 and FLVCR2 are localized at the plasma membrane (arrows). Plasma membrane GFP (mGFP) was used as a marker. **b**, Choline transport activities of human FLVCR1 and FLVCR2. Choline kinase A (*CHKA*) was co-expressed with both proteins. **b**,**c**, Dose curves and time courses for choline transport activities of human FLVCR1 and FLVCR2. *CHKA* was co-expressed with both proteins. **d**, Ethanolamine transport activities of human FLVCR1 and FLVCR2. Ethanolamine kinase 1 (*ETNK-1*) was co-expressed with both proteins. **e**,**f**, Dose curves and time courses for ethanolamine transport activities of human FLVCR1 and FLVCR2. *ETNK-1* was co-expressed with both proteins. The inactive S203Y mutant of FLVCR2 was used as a control in all experiments. Each symbol represents one replicate. Data are expressed as mean ± SD. ****P<0.0001, ***P<0.001. ns, not significant. One-way ANOVA for transport activity measurement; Two-way ANOVA for dose curve measurements.

### FLVCR architecture and conformational landscape

Next, we aimed to establish structure-function relationships for the choline and ethanolamine transport properties of FLVCR1 and FLVCR2. Wildtype FLVCR1 and FLVCR2 proteins were purified and subjected to SPA cryo-EM (Supplementary Fig. 1a-f). We determined inward-facing conformation structures of FLVCR1 and FLVCR2 in their apo states at 2.9 Å resolution (Fig. 2a,b, and Supplementary Fig. 2,3). In addition, we successfully elucidated the outward-facing conformational structure of FLVCR2 at 3.1 Å resolution from the same sample preparation (Fig. 2c and Supplementary Fig. 3). Both FLVCR paralogs share a common MFS-type architecture^28^, with their N-domain containing transmembrane domain (TMs) 1-6 and the C-domain containing TMs 7-12 connected by a long and flexible loop containing two short horizontal cytoplasmic helices (H1 and H2) (Extended Data Fig. 4a,b). In the FLVCR2 structure, we resolved a short helical segment at the C-terminus (H3), while the density of the FLVCR1 C-terminus was less pronounced (Extended Data Fig. 4a,b and Supplementary Fig. 4). Further, we identified an N-linked glycosylation site at N265 of FLVCR1 which locates within the extracellular loop connecting TMs 5 and 6 (EL5-6) (Fig. 2a). In contrast, no glycosylation site was identified in the FLVCR2 structure^19^ (Fig. 2b,c and Supplementary Fig. 5).

**Fig. 2:**
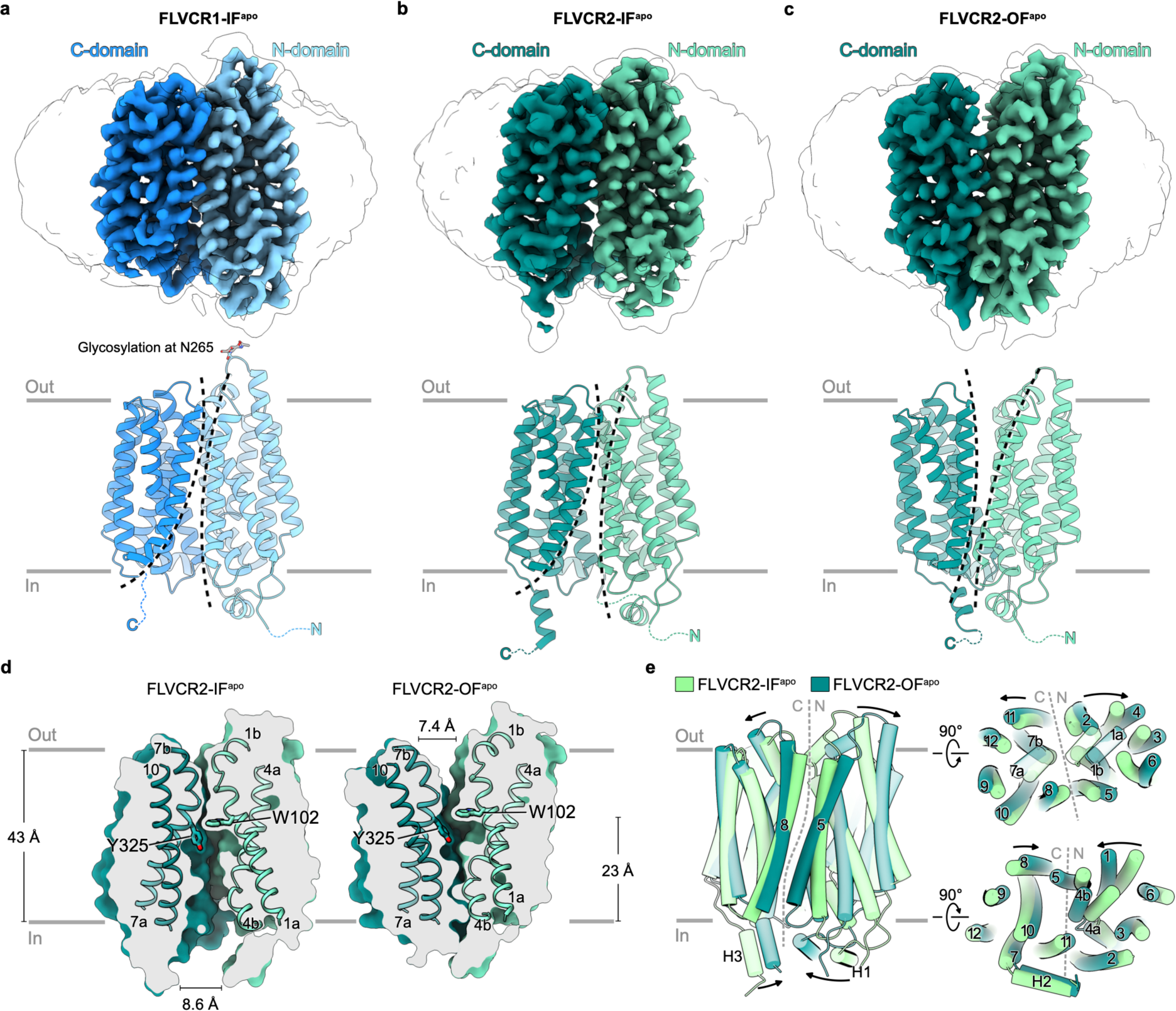
Architecture of FLVCR1 and FLVCR2 in their inward- and outward-facing states. Cryo-EM density (top) and atomic model (middle and bottom) of apo inward-facing FLVCR1 (**a**) as well as FLVCR2 in the inward-facing (**b**) and outward-facing (**c**) states. The N- and C-domains are coloured in different shades of blue and green for FLVCR1 and FLVCR2, respectively. A transparent cryo-EM density lowpass-filtered at 6 Å is shown to visualize the detergent belt surrounding the transmembrane region. The sidechain of N265 in FLVCR1 and its N-linked glycan (grey) are shown as stick models. **d**, Cut-away views of the surface representation showing the cavity shape of FLVCR2-IF^apo^ (left) and FLVCR2-OF^apo^ (right). Two central aromatic residues are shown as sticks. **e**, Structural superposition of FLVCR2-OF^apo^ (dark green) and FLVCR2-IF^apo^ (light green).

The inward-facing conformations of FLVCR1 and FLVCR2 exhibit a close resemblance to each other, with a root mean square deviation (C*α* r.m.s.d.) of 0.993 Å (Extended Data Fig. 4b). Both structures feature a wedge-shaped solvent-accessible cavity that is mainly created by the separation of TMs 4 and 5 of the N-domain from TMs 10 and 11 of the C-domain (Fig. 2d and Extended Data Fig. 4b and 5a). This space extends halfway across the membrane with a similar depth of around 23 Å in both transporters. On the level of the outer leaflet, TMs 1, 2, and 5 of the N-domain and TMs 7, 8, and 11 of the C-domain pack tightly against each other and thus shield the central cavity from the extracellular space (Fig. 2d and Extended Data Fig. 4b and 5a). Notably, TMs 1, 4, and 7 exhibit disordered regions at their C-terminal ends, followed by short kinked helical motifs designated TMs 1b, 4b, and 7b (Fig. 2d and Extended Data Fig. 4a and 5a).

The outward-facing conformation of FLVCR2 features a cavity accessible from the extracellular space. This cavity is formed through a ‘rocker-switch’ rigid-body motion that occurs during the transition from the inward- to outward-facing state (Fig. 2d, e and Supplementary Video 1). This motion involves a shift of the outer halves of all transmembrane segments away from the central axis. Concurrently, the inner halves of these TMs move inward, effectively blocking the exit pathway. (Fig. 2e). The cavities in outward- and inward-facing conformations are 7.4 Å and 8.6 Å wide at their respective openings (Fig. 2d). Both inward- and outward-facing cavities are lined by uncharged and hydrophobic residues which are mostly conserved (Extended Data Fig. 6). Compared to the inward-facing cavity, the outward-facing cavity demonstrates a more restricted pathway within its central region. In this narrowed segment of the cavity, the channel ends at W102^FLVCR2^ and Y325^FLVCR2^ (equivalent to W125^FLVCR1^ and Y349^FLVCR1^), two residues that are highly conserved in both FLVCR transporters (Fig. 2d and Extended Data Fig. 5a and 6).

In line with an alternating-access model, the inward-facing conformation of FLVCR2 features a tightly-sealed extracellular gate created by inter-domain interactions. This is mainly achieved by the juxtaposition of TM1b (N-domain) and TM7b (C-domain) (Fig. 2d and Extended Data Fig. 4b). The inter-domain interaction between these two motifs is stabilized by a hydrogen bonding network consisting of two pseudo-symmetry-related asparagine residues N110^FLVCR2^ and N332^FLVCR2^, and E343^FLVCR2^ (TM8) and N239^FLVCR2^ (EL5-6) (Extended Data Fig. 5a and Supplementary Fig. 6a). Furthermore, we identified a stable inter-domain salt bridge between D124^FLVCR2^ (TM2) and R333^FLVCR2^ (TM7b), reinforcing the external occlusion (Extended Data Fig. 5a and Supplementary Fig. 6b). Our cell-based mutagenesis studies show that alanine substitutions at N110^FLVCR2^, E343^FLVCR2^, D124A^FLVCR2^ and R333A^FLVCR2^ individually result in a significant decrease of choline uptake and an almost complete perturbation of ethanolamine transport, highlighting a critical role of inter-domain interactions on the extracellular surface for FLVCR2 functionality (Extended Data Fig. 5b and Supplementary Fig. 7). While FLVCR1 exhibits a similar extracellular hydrogen bonding network as FLVCR2, it lacks a corresponding salt bridge interaction (Extended Data Fig. 5a). Notably, the disruption of the hydrogen bond between N133^FLVCR1^ and N359^FLVCR1^ (structurally analogous to N110^FLVCR2^ and N332^FLVCR2^) through an alanine substitution of N133^FLVCR1^ markedly diminishes FLVCR1’s transport activity for both choline and ethanolamine. In contrast, the E367A^FLVCR1^ mutation impacts the transport of choline to a stronger extend as compare to ethanolamine transport (Extended Data Fig. 5c).

During the transition from inward- to outward-facing conformation of FLVCR2, inter-domain interactions contributing to the extracellular gate become disrupted, while the formation of the intracellular gate occludes the central cavity from the cytoplasmic side. TM4b of the N-domain moves in proximity to the N-terminal end of TM11 of the C-domain to establish a first level of occlusion (Fig. 2d,e). An interaction network consisting of several hydrogen bonds and a salt bridge is found within this region as well (Extended Data Fig. 5a). The E435^FLVCR2^ residue in TM11 plays a pivotal and multifaceted role, forming a hydrogen bond with S203^FLVCR2^ and simultaneously establishing a salt bridge with R200^FLVCR2^ in TM4b (Extended Data Fig. 5a and Supplementary Fig. 6b). Individuals with a homozygous S203Y mutation in FLVCR2 are nonviable, likely attributable to the complete loss of choline and ethanolamine transport activity^29^ (Fig. 1b,c). Our mutagenesis studies underscore a greater significance of S203^FLVCR2^ compared to R200^FLVCR2^, particularly in choline transport (Fig. 1b and Extended Data Fig. 5b). This suggests that the interaction between E435^FLVCR2^ and S203^FLVCR2^ is essential for either facilitating conformational changes during the transport cycle or maintaining the stability of FLVCR2 in its outward-facing state. An additional inter-domain hydrogen bond is identified between S199^FLVCR2^ in TM4b and S439^FLVCR2^ in TM1 (Extended Data Fig. 5a and Supplementary Fig. 6a). In the peripheral region, K372^FLVCR2^ and R374^FLVCR2^ (EL8-9) approach N209^FLVCR2^ (EL4-5) and S212^FLVCR2^ (TM5) to form hydrogen bond pairs and thus block the lateral accessibility of the cavity (Extended Data Fig. 5a and Supplementary Fig. 6a). A second level of occlusion was observed beneath the cytoplasmic ends of TMs 4, 10 and 11, where H1 and H3 are positioned in close proximity (Extended Data Fig. 5a). Here, the backbone carbonyl group and amide nitrogen of A283^FLVCR2^ (H1) form stable hydrogen bonds with N497^FLVCR2^ (H3) and Y431^FLVCR2^ (IL10-11), respectively (Extended Data Fig. 5a and Supplementary Fig. 6a). Together with the loop connecting TMs 10 and 11, the two helical motifs H1 and H3 serve as a cytoplasmic latch to secure the closure of the two domains.

### Choline binding and coordination chemistry of FLVCR1 and FLVCR2

To decipher the substrate binding and coordination chemistry of FLVCR1 and FLVCR2, we employed choline as a ligand and determined their structures at resolutions of 2.6 Å and 2.8 Å, respectively (Fig. 3a,b and Supplementary Fig. 2, 3 and 4d,e). Both structures were captured in the inward-facing conformation, suggesting that choline is capable of resolving the previously captured conformational heterogeneity of FLVCR2. The choline-bound structures exhibit a C*α* r.m.s.d. of 0.688 Å (FLVCR1) and 1.002 Å (FLVCR2), respectively, compared to their apo inward-facing structures (Extended Data Fig. 7a,b). In the choline-bound structures, the binding sites are located at analogous positions, with the choline molecule situated between the two domains, surrounded mainly by TMs 1, 2, 4, 5, 7, and 11 (Fig. 3a,b). The binding site in both transporters comprises an identical composition of conserved residues. We observed that W102^FLVCR1^ and W125^FLVCR2^ in TM1 directly interact with choline. This central coordinating tryptophan residue is located above choline, protruding its side chain to constrain the diffusion of the molecule towards the extracellular space (Fig. 2d and Fig 3c,d). Two additional aromatic residues of TM7, one tyrosine (Y325^FLVCR1^ and Y349^FLVCR2^) and one phenylalanine (F348^FLVCR1^ and F324^FLVCR2^), line the peripheral space of the binding site and restrict the movement of choline within the pocket. Our MD simulations demonstrate that the quaternary ammonium group of choline forms stable cation-π interactions with the conserved tryptophan (W125^FLVCR1^ and W102^FLVCR2^) and tyrosine residues (Y325^FLVCR1^ and Y349^FLVCR2^) in a simultaneous manner (Fig. 3c,d). Mutations of W125A^FLVCR1^ and W102A^FLVCR2^ significantly reduced choline transport activity (Fig. 3e and Supplementary Fig. 7). MD simulations further confirmed the key role of the tryptophan in both transporters (Fig. 3f,g). We also found that the hydroxyl group of choline is essential for ligand recognition by both FLVCRs. Substitution to a carboxylic group as found in betaine abolishes ligand binding and transport activity^26^ (Fig. 3h). In our choline simulations, this hydroxyl shows versatile interactions by forming transient hydrogen bonds with two proximal asparagine residues (Q214^FLVCR1^/Q471^FLVCR1^, Q191^FLVCR2^/Q447^FLVCR2^). Further interactions come from at least two water molecules that were consistently present around the hydroxyl moiety of the choline (Supplementary Fig. 8 and Supplementary Video 2). Functional analyses of Q214^FLVCR1^, Q471^FLVCR1^, Q191^FLVCR2^, and Q447^FLVCR2^ indicate that changes of the local protein environment around the hydroxyl group of choline, caused by alanine substitutions, discernibly affect the transport activities of both FLVCRs (Fig. 3e). It is noteworthy that all these residues are fully conserved in mammalian FLVCR homologs, suggesting a common substrate profile of FLVCRs across species (Fig. 3i).

**Fig. 3:**
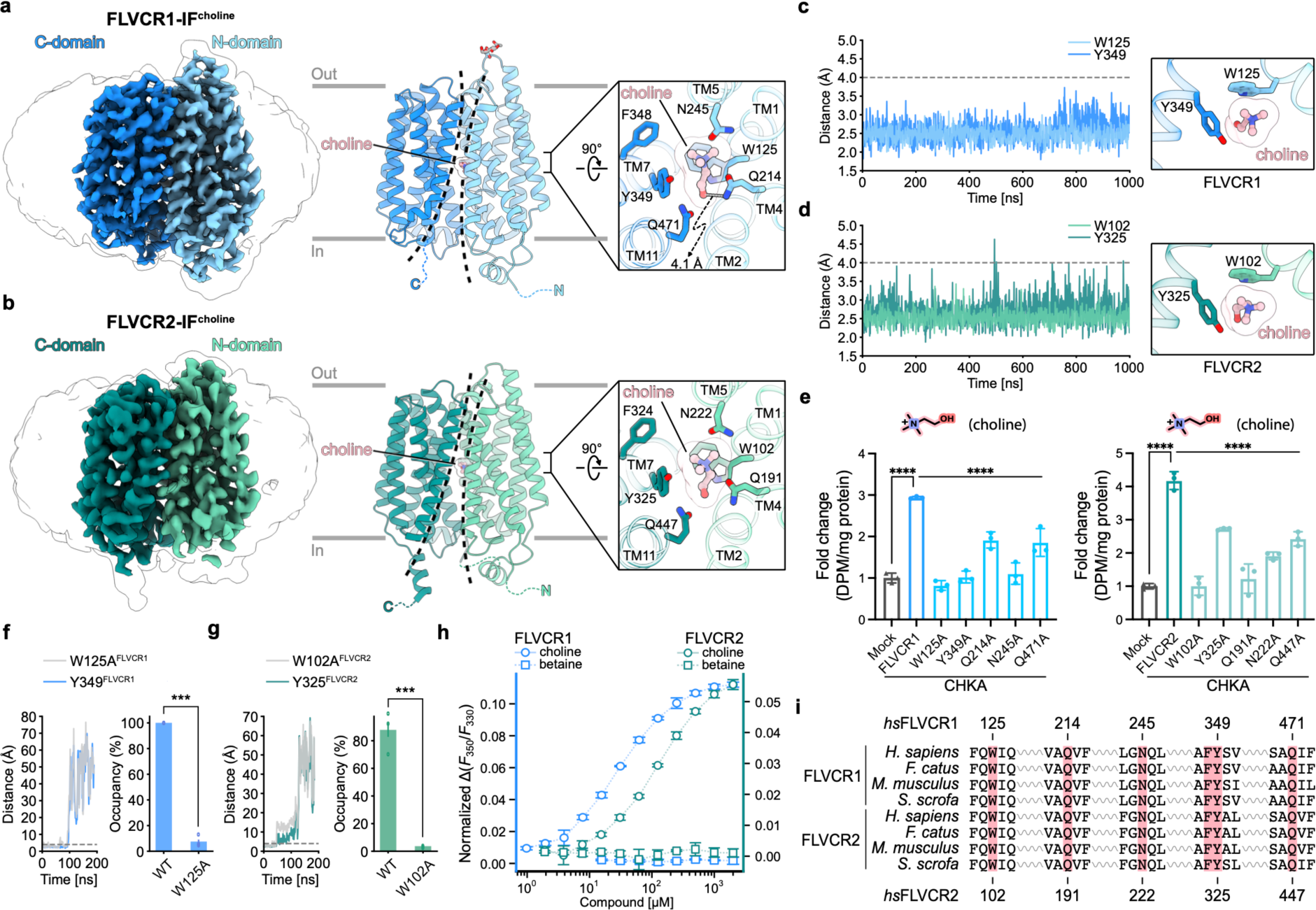
Cryo-EM structures of FLVCR1 and FLVCR2 in complex with choline. **a**,**b**, Cryo-EM densities and atomic models of the choline-bound inward-facing FLVCR1 (FLVCR1-IF^choline^) structure (**a**), and the choline-bound inward-facing FLVCR2 (FLVCR2-IF^choline^) structure (**b**), respectively. The bound choline is shown as ball-and-stick model; binding site residues are shown as sticks. **c**,**d**, Time-resolved distance plots between choline and the highly-conserved tryptophan and tyrosine residues forming the choline-binding pockets of FLVCR1 (**c**) and FLVCR2 (**d**) obtained from MD simulation runs. A cation-π interaction is assumed for distance < 4 Å (grey dashed line). **e**, Choline transport activity of indicated FLVCR1 and FLVCR2 mutants. *CHKA* was co-expressed with all proteins. 20 µM [^3^H] choline was used for FLVCR1 mutants and 100 µM [^3^H] choline was used FLVCR2 mutants, respectively. Each symbol represents one replicate. Mutant transport activities were normalized to the total protein from cells and displayed as fold change with reference to mock (empty vector). **f**,**g**, Distance of choline to conserved aromatics for W125A^FLVCR1^ (**f**) and W102A^FLVCR2^ (**g**) as function of time in MD simulations (left), and choline occupancy in binding site for WT and alanine mutants (right). Error bars represent standard errors of the mean (s.e.m.) **h**, Shifts of tryptophan fluorescence of FLVCR1 and FLVCR2 in the presence of choline or betaine. **i**, Protein sequence alignment of choline-binding pocket residues (red block) in FLVCR1 and FLVCR2 across various mammalian species. Indicated residue numbers refer to FLVCR1 and FLVCR2 from *homo sapiens*. Data shown are mean ± SD for (**e**) and (**h**), and mean ± standard error of the mean (s.e.m) for (**f**). ****P<0.0001, ***P<0.001. One-way ANOVA for (**e**), and *t*-test for (**f**) and (**g**).

### Molecular mechanism of ligand selectivity of FLVCR1

Guided by our functional analyses, we advanced our investigation into the molecular mechanism of ethanolamine binding by FLVCR1. We obtained the cryo-EM structure of FLVCR1 complexed with ethanolamine at a resolution of 3.1 Å, showing an inward-facing conformation analogous to the choline-bound FLVCR1 structure (Fig. 4a, Extended Data Fig. 7c and Supplementary Fig. 2). The ethanolamine density was identified within the abovementioned ligand binding-pocket, with its primary amine group engaging in cation-π interactions with the conserved W125^FLVCR1^ and Y349^FLVCR1^ residues (Fig. 4b, and Supplementary Fig. 4f). Our cryo-EM data, corroborated by MD simulations, indicate that ethanolamine is positioned slightly further from W125^FLVCR1^ and Y349^FLVCR1^ than choline, attributable to differences in their molecular orientations (Fig. 3a, 4a and 4c, and Extended Data Fig. 8). While the quaternary ammonium group of choline is situated near Q214^FLVCR1^, ethanolamine’s orientation results in its amine group being distanced, consequently bringing its hydroxyl group into closer proximity to Q214^FLVCR1^ (Fig. 4d). These nuanced molecular differences presumably contribute to the selective substrate profile of FLVCR1 as observed in our cell-based transport assays where FLVCR1 showed a preferential ethanolamine transport activity (Fig. 1b, c, 3e and 4e). This is further substantiated by the Q214A^FLVCR1^ mutant variant, which results in a complete loss of ethanolamine transport activity while only partially affecting choline transport (Fig. 3e and 4e). Notably, alanine substitution of the analogous Q191^FLVCR2^ residue abolishes the transport of both ethanolamine and choline in FLVCR2 (Fig. 3e and 4e).

**Fig. 4:**
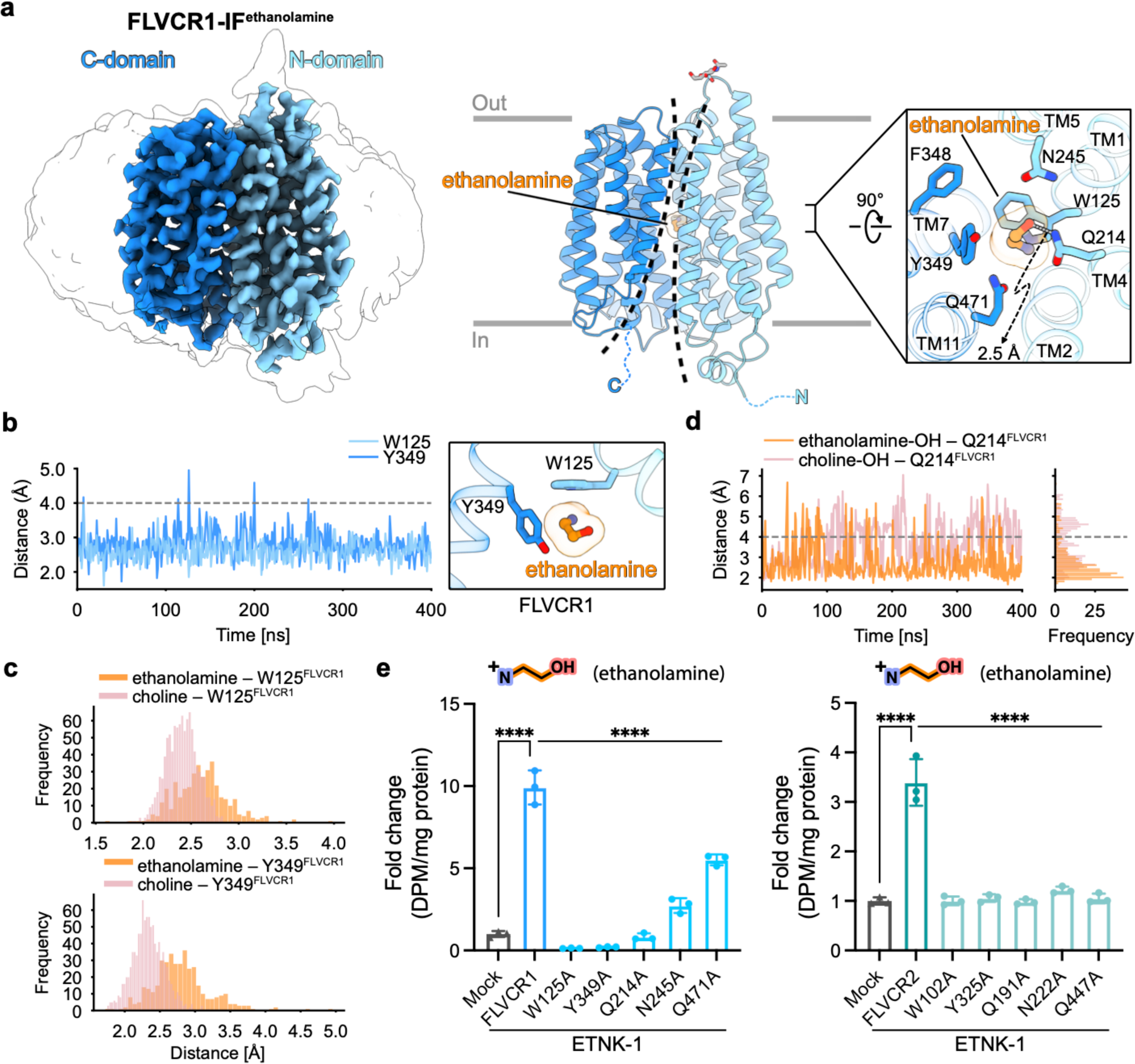
Cryo-EM structures of FLVCR1 and FLVCR2 in complex with ethanolamine. **a**, Cryo-EM density and atomic model of the ethanolamine-bound inward-facing FLVCR1 (FLVCR1-IF^ethanolamine^) structure. The bound ethanolamine is shown as ball-and-stick model; binding site residues are shown as sticks. **b**, Distance of ethanolamine to conserved aromatics of FLVCR1 as function of time in MD simulation (left) and snapshot (right). **c**, Frequency of distance between ethanolamine or choline and W125^FLVCR1^ (top) or Y349^FLVCR1^ (bottom) derived from Fig. 3c and 4b, respectively. **d**, Distance between Q214^FLVCR1^ and the hydroxyl group of ethanolamine (orange) or choline (pink) as function of time in MD simulation, with corresponding frequency (right). **e**, Ethanolamine transport activities of indicated FLVCR1 and FLVCR2 mutants. *ETNK-1* was co-expressed with all proteins. 2.5 µM [^14^C] ethanolamine was used. Transport activities of mutant variants were normalized to the total protein from cells and are displayed as fold change to mock (empty vector). Each symbol represents one replicate. Data shown are mean ± SD. ****P<0.0001. One-way ANOVA.

### Identification of a peripheral heme binding site of FLVCR2

Our cell-based assays, using hemin as a heme analog substrate, revealed that unlike the choline and ethanolamine uptake data, neither FLVCR1 nor FLVCR2 demonstrated heme import activity (Fig. 5a). This observation was confirmed by our structural studies which did not reveal any density for bound heme molecules within the ligand binding cavities of FLVCR1 or FLVCR2, respectively (Fig. 5b). Of note, neither the dimensions, nor the residues lining the central cavities of both FLVCRs indicate a structural adaptation for accommodating a heme molecule as observed in other heme transporters^30–32^.

**Fig. 5:**
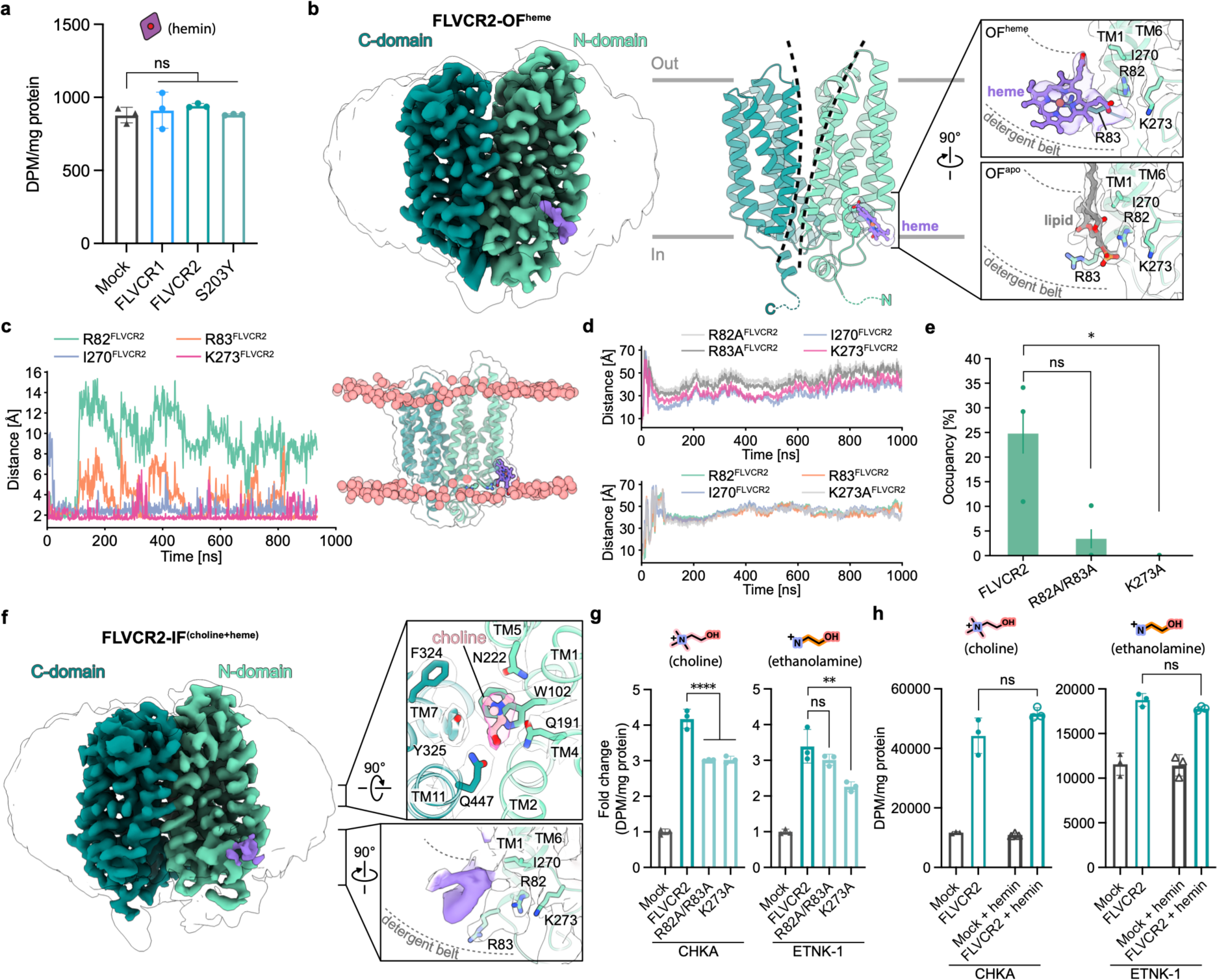
Cryo-EM structures of FLVCR2 in complex with heme. **a**, Import assay of FLVCR1 and FLVCR2 for heme. 2.5 µM [^3^H] hemin was used. The inactive mutant of FLVCR2^S203Y^ was used as a control. **b**, Cryo-EM density (left) and ribbon model (right) of the heme-bound outward-facing FLVCR2 (FLVCR2-OF^heme^). The heme molecule is shown in ball-and-stick representation, and the corresponding cryo-EM density is coloured in purple. Right panels are the close-up views of the heme-binding site in FLVCR2-OF^heme^ (top) and in FLVCR2-OF^apo^ (bottom) with locally-filtered cryo-EM density. Residues in close proximity to heme are shown as sticks. A lipid molecule (grey ball-and-stick) is fitted into the lipid-like density at the heme-binding site of FLVCR2-OF^apo^. **c**, Distance of heme and interacting residues as function of time in MD simulations (left), with a snapshot of FLVCR2-heme interactions (right) indicating the membrane layer by the phosphates of POPE/POPG (red spheres). **d**, Distances as in (**c**, left), but for mutants R82A/R83A (top) and K273A (bottom). **e**, Interaction frequency of heme with four heme-interacting residues for WT and mutants of FLVCR2. **f**, Cryo-EM density of the heme-bound, choline-bound, inward-facing FLVCR2. The cryo-EM density corresponding to heme is coloured in purple. Right panels are the cryo-EM density of the substrate-binding site (top) and heme-binding site (bottom) in FLVCR2-IF^heme-choline^ with rigid-body fitting of the FLVCR2-IF^choline^ model. The heme-binding site is shown with locally-filtered cryo-EM density. **g**, Choline (left) and ethanolamine (right) transport assays for *FLVCR2* heme-binding site mutants. *CHKA* or *ETNK-1* was co-expressed with WT or *FLVCR2* mutant plasmids. 100 µM [^3^H] choline or 2.5 µM [^14^C] ethanolamine was used, respectively. Transport activity of the mutants was normalized to the total protein and is displayed as fold change to mock (empty vector). **h**, Choline (left) and ethanolamine (right) transport assay for FLVCR2 in the presence of heme. For choline + hemin experiment, 10 µM [^3^H] choline and 50 µM hemin were used, while for ethanolamine + hemin experiment, 2.5 µM [^14^C] ethanolamine and 25 µM hemin were used. The experiment was repeated twice for (**a**). One dataset was shown. Each symbol represents one replicate. Data are expressed as mean ± SD for (**a**), (**g**) and (**h**), and mean ± s.e.m for (**e**). ****P<0.0001, **P<0.01, *P<0.05. ns, not significant. One-way ANOVA for (**a**) and (**g**), two-way ANOVA for (**h**), and *t*-test for (**e**).

However, upon determining the structure of FLVCR2 in the presence of heme we noticed that heme resolved the previously characterized conformational heterogeneity, fully driving the transporter into the outward-facing conformation (Fig. 5b and Supplementary Fig. 9). Additionally, we identified a density near the N-terminus of TM1 on the intracellular side of FLVCR2, likely corresponding to heme. This density appears to have replaced a peripherally bound lipid molecule that we had previously observed in the outward-facing density map of apo FLVCR2 (Fig. 5b). The protein surface in this particular region displays a patch of positive charges, suggesting the heme molecule to be held in place by the formation of electrostatic interactions between its two propionate groups and the side chains of R81^FLVCR2^, R82^FLVCR2^, and K273^FLVCR2^ (Fig. 5b and Extended Data Fig. 6). The map density of bound heme in our structure indicates local mobility of the heme, possibly due to the absence of axial coordination of the central iron in the macrocycle scaffold (Fig. 5b). By contrast, cryo-EM studies on FLVCR1 in the presence of heme did not reveal any heme binding or notable conformational changes.

To further investigate heme binding in FLVCR2, MD simulations were performed within a lipid bilayer. Heme was found to interact with the intracellular loop region rather than accessing the central cavity, in agreement with prior functional and structural observations (Supplementary Video 3). When placed near the binding site at the N-terminus of TM1 as indicated by our cryo-EM data, heme rapidly disengaged from R82^FLVCR2^, yet maintained contact with R83^FLVCR2^, K273^FLVCR2^, and I270^FLVCR2^ after insertion into the lipid bilayer (Fig. 5c,d and Supplementary Fig. 10). Subsequent simulations with alanine substitution variants of these residues led to either reduction or complete loss of heme-binding events, indicating the involvement of these residues for FLVCR2 interaction with heme (Fig. 5e).

Follow-up structural studies conducted in a competitive manner by the presence of both heme and choline revealed that choline binding occurs within the central cavity of FLVCR2 irrespective of the presence of heme and triggers conformational shifts towards the inward-facing state. This occurs despite heme’s ability to drive FLVCR2 into an outward-facing conformation in the absence of choline (Fig. 5f and Supplementary Fig. 9). Our results from the cell-based transport assays showed that the introduction of mutations at the heme-binding residues marginally influenced the choline and ethanolamine transport function of FLVCR2 (Fig. 5g). Nevertheless, co-incubation of heme with either choline or ethanolamine did not impact the transport activity of these ligands in FLVCR2 (Fig. 5h). In conclusion, our structural studies reveal a peripheral heme binding site in FLVCR2 that does not play a critical role in choline and ethanolamine transport.

### Translocation pathway of choline and ethanolamine in FLVCRs

MFS transporters typically cycle between inward- and outward-facing conformations, facilitating substrate translocation in an alternating-access manner, and substrate binding plays a pivotal role in eliciting the conformational transitions^33^. Our cryo-EM data supports this mechanism in FLVCR2, where we see a full transition from the outward-facing to inward-facing state upon choline binding. Complementary to our structural insights into the conformational landscape, we performed MD simulations to map the route for choline entry along the pathway in the outward-facing conformation of FLVCR2. After spontaneously diffusing into the translocation pathway, choline initially interacts with several residues near the protein surface, mainly D124^FLVCR2^ (Supplementary Fig. 11). It sequentially approaches the deeper recesses of the cavity, primarily engaging with conserved aromatic residues W102^FLVCR2^ and Y325^FLVCR2^. In one of the entry events, we observed choline moving spontaneously to a position below the W102^FLVCR2^ residue within the binding site consistent with our structural data (Supplementary Video 4).

The substantial global conformational changes triggered by substrate further alter the local arrangement of substrate-coordinating residues within the translocation pathway as seen in our cryo-EM maps. The rearrangement of the binding site results in a more constricted pocket, promoted by the inward movement of the conserved residues towards the substrate molecule (Extended Data Fig. 9a). The repositioning of these residues, especially the conserved aromatic side chains W102^FLVCR2^, F324^FLVCR2^, and Y325^FLVCR2^, (equivalent to W125^FLVCR1^, F348F^LVCR1^, and Y349^FLVCR1^) restricts the accessibility of the binding pocket and promotes substrate engulfment and coordination. In their choline-bound structures, the cavities of both FLVCRs share common characteristics, exhibiting a neutral interior but a negatively charged surface at the exit (Extended Data Fig. 6b, e). FLVCR1 features a slightly smaller cavity volume of 513 Å^3^ compared to 579 Å^3^ of FLVCR2 (Extended Data Fig. 9b). While the cavity narrows toward the intracellular side, a peripheral solvent-accessible channel emerging from the cytoplasmic space to the binding site reveals a semi-open translocation pathway in the substrate-bound inward-facing conformations of both transporters, which may facilitate the release of the bound ligand (Extended Data Fig. 9b).

## Discussion

While FLVCR1 has many studies linking its activity to heme export^8–12^, the study of FLVCR2-mediated heme uptake has not been confirmed^13^, and furthermore, conclusive biochemical and detailed molecular evidence had remained elusive for the function of both transporters^14–17^. In this study, we have investigated the choline and ethanolamine transport properties of FLVCR1 and FLVCR2 via cell-based transport assays and determined cryo-EM structures of human FLVCR variants in distinct apo, choline-bound, and ethanolamine-bound states. Our work provides valuable insights into architecture, ligand binding-chemistries, and conformational landscapes of these two MFS transporters. Via multiple lines of evidence, we arrive at the conclusion that choline and ethanolamine represent primary transport substrates of both FLVCR1 and FLVCR2, while also revealing a peripheral heme-binding site of FLVCR2 with yet unknown functional implications^10,16^. Our findings suggest that both FLVCR1 and FLVCR2 operate as uniporters, facilitating downhill ligand transport independent of sodium or pH gradients. Notably, FLVCR1 and FLVCR2 possess similar ligand binding and coordination chemistries in which a key conserved tryptophan residue (W125A^FLVCR1^ /W102A^FLVCR2^) forms a cation-π interaction with the ammonium and amine groups of the respective ligands. Further, we characterized the functional importance of several glutamine residues within the substrate-binding site that participate in hydrogen bonds with the hydroxyl groups of choline and ethanolamine during ligand coordination or transport. Functional assays point to a ligand preference between FLVCR1 and FLVCR2. Our structural studies suggest that ligand coordination is a crucial determinant of the transport preference between choline and ethanolamine for both FLVCRs. Given that most solute carrier transporters exhibit ligand promiscuity^34,35^, the specific role of FLVCR1 as an ethanolamine transporter and FLVCR2 as a choline transporter in physiological conditions has yet to be confirmed through *in vivo* studies in animal models.

Based on our structural findings and simulation data, we suggest a rocker-switch alternating-access mechanism for the transport cycle of choline import (Fig. 6, and Supplementary Video 4 and 5). We propose a transport cycle model for both FLVCRs in which the outward-facing conformation represents the state for ligand binding from the extracellular space. Substrate-induced conformational changes drive the transporter towards its inward-facing state. Finally, ligands are released to the intracellular space to enter metabolic pathways. Subsequent to choline-release, FLVCRs will undergo additional conformation changes to adopt their outward-facing state in order to re-initiate the transport cycle. Our structural data did not reveal fully occluded conformations that are key features of alternating access. Hence, we suspect that these occluded conformations exist only transiently and rapidly convert towards either the inward- or outward-facing conformations^33,36,37^.

**Fig. 6:**
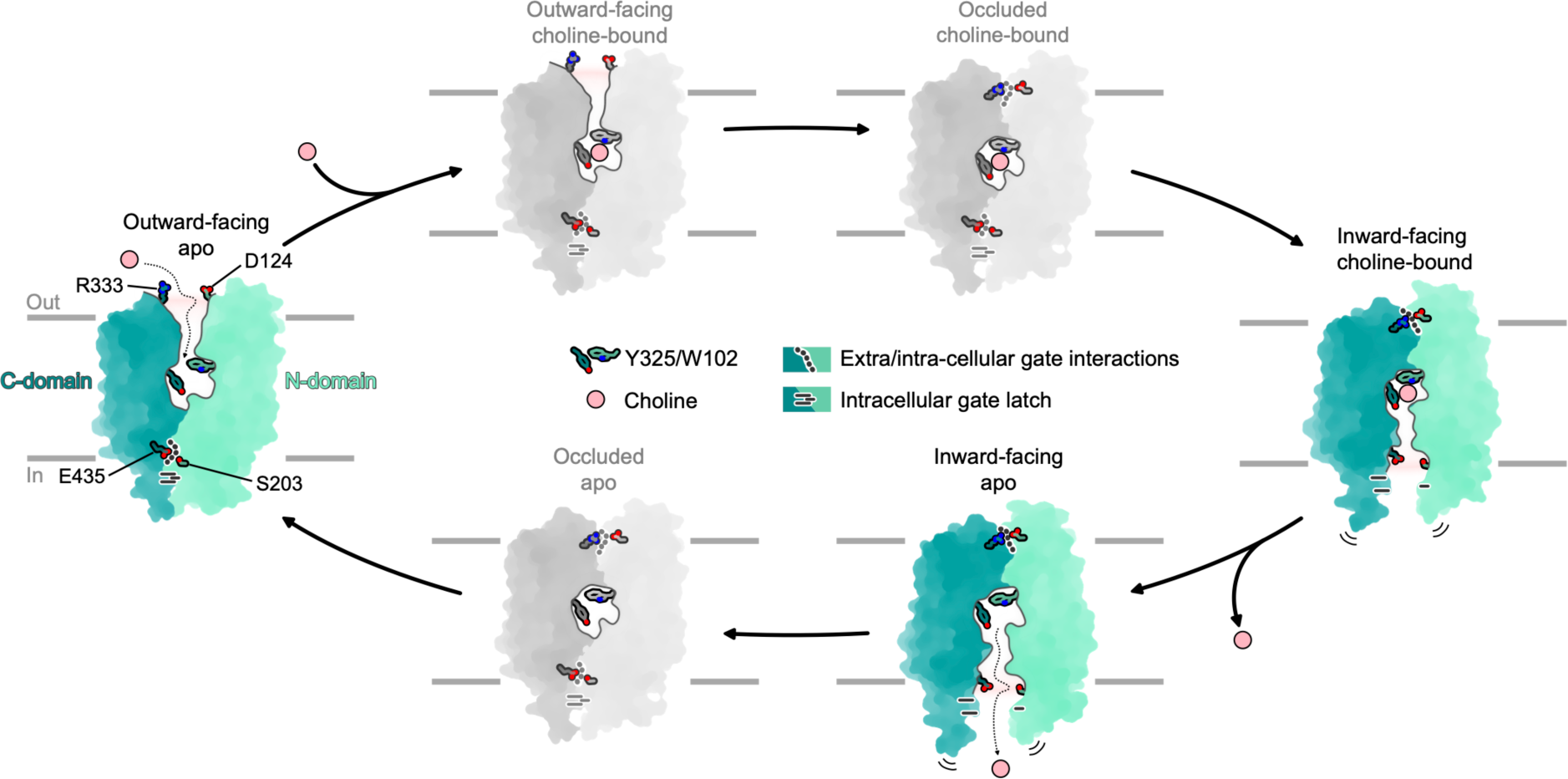
Proposed model for choline transport by FLVCR2. Schematic illustration of FLVCR2 conformations during the choline transport cycle. Green-colored states represent experimentally obtained conformations of this study. States colored in grey are hypothesized based on knowledge about the commonly characterized alternative-access mechanism of MFS transporters.

In summary, our research delineates the structural framework through which FLVCR1 and FLVCR2 mediate the cellular transport of choline and ethanolamine. Identifying these proteins as facilitative transporters provides a crucial groundwork for future characterizations of the physiological functions of choline and ethanolamine transport *in vivo*. Furthermore, our findings provide mechanistic insights for understanding of the disease-mechanisms pertaining to the mutations of these essential genes.

## Online Methods

### Generation of inducible HEK293 stable cell lines

The complementary DNAs of full-length wildtype FLVCR1 (human SLC49A1, NCBI Reference Sequence NM_014053) and FLVCR2 (human SLC49A2, NCBI Reference Sequence: NM_017791) were cloned into pcDNA5/FRT/TO (Invitrogen) vectors, respectively. The gene for both FLVCRs was modified by a C-terminal FLAG fusion tag. Further details are found in sequence data provided as supplementary information (Supplementary Tables 2 and 3). The recombinant Flp-In T-REx293-FLVCR1 and Flp-In T-REx293-FLVCR2 cell lines were generated by using a tetracycline-inducible and commercially available Flp-In T-REx host-cell line system from Invitrogen. Flp-In T-REx293 cells were cultured in high-glucose Dulbecco’s Modified Eagle’s Medium (DMEM; Sigma-Aldrich) supplemented with 10% fetal bovine serum (FBS; Gibco), 1% Pen/Strep (Gibco), 1 μg ml^-1^ Zeocin (Thermo Fisher), and 15 μg ml^-1^ blasticidin S hydrochloride (AppliChem) at 37 °C in an atmosphere of 5% CO_2_. Cells were not tested for mycoplasma contamination. For stable integration, the pcDNA5/FRT-FLVCR1-FLAG and pcDNA5/FRT-FLVCR2-FLAG vectors were cotransfected with the Flp recombinase encoding expression vector pOG44 (Thermo Fisher) at a 1:13 mass ratio, respectively. All transfection procedures were performed with Lipofectamine™ 2000 reagent according to the manufacturer’s instructions (Thermo Fisher). To select for stable clones, transfected cells were cultivated with growth medium containing 100 μg ml^-1^ hygromycin B (AppliChem).

### Transport assays in HEK293 cells

HEK293 cells were co-transfected with pcDNA3.1 plasmid and human *FLVCR1* or *FLVCR2* and human choline kinase A (*CHKA*) for choline transport assays or ethanolamine kinase 1 (*ETNK-1*) for ethanolamine transport assays using Lipofectamine™ 2000 reagent (Thermo Fisher Scientific). After 24 hours post-transfection, cells were incubated with DMEM containing 20 μM [^3^H] choline or 2.5 μM [^14^C] ethanolamine. The cells were incubated at 37 °C and 5% CO_2_ for 1 hour for uptake of the ligands. The cells were subsequently washed with ice-cold plain DMEM and lysed with RIPA buffer by shaking for 30 minutes. The cell lysates were also quantified by scintillation counter Tricarb. Radioactive signals from cell lysates were normalized to total protein levels. For dose curve assays, indicated concentrations of choline and ethanolamine were incubated with the cells for 1 hour at 37 °C. For time course assays, the cells were incubated with 20 μM [^3^H] choline or with 2.5 μM [^14^C] ethanolamine. The transport assays were stopped at indicated time points by adding ice-cold plain DMEM. For testing transport activity of FLVCR1 mutants, 20 μM [^3^H] choline and 2.5 μM [^14^C] ethanolamine was used. For testing transport activity of FLVCR2 mutants, 100 μM [^3^H] choline and 2.5 μM [^14^C] ethanolamine was used. For transport assays of HEK293 cells overexpressed with *FLVCR1* or *FLVCR2* without co-expressing with *CHKA* or *ETNK-1*, 20 μM [^3^H] choline and 2.5 μM [^14^C] ethanolamine was used.

For transport assays with indicated pH conditions, the following buffers were used: pH8.5 buffer (140 mM NaCl, 20 mM Tris-HCl pH 8.5, 2 mM CaCl_2_, 1 g/L D-glucose), pH6.5 buffer (140 mM NaCl, 20 mM MES pH 6.5, 2 mM CaCl_2_, 1 g/L D-glucose), or pH7.5 buffer (140mM NaCl, 20mM HEPES-NaOH, 2mM CaCl2, 1 g/L D-glucose). For sodium-free buffer, buffer containing 140 mM KCl, 20 mM HEPES-KOH pH 7.5, 2 mM CaCl_2_, 1 g/L D-glucose was used. In these assays, 20 μM [^3^H] choline or 2.5 μM [^14^C] ethanolamine was used, and the assays were stopped after 15 minutes of incubation with the ligands. Radioactive signals from cell lysates were normalized to total protein levels.

### Immunofluorescent staining (IF)

HEK293 cells were seeded onto 24-well plates with coverslips and maintained in Dulbecco’s Modified Eagle’s Medium (Gibco) supplemented with 10% fetal bovine serum and 1% penicillin-streptomycin. HEK293 cells were co-transfected with *FLVCR1* or *FLVCR2* with membrane expressing GFP (Addgene: #14757) using Lipofectamine™ 2000 reagent (Thermo Fisher Scientific). The inducible HEK293 stable cell lines overproducing FLVCR1 or FLVCR2 were seeded onto Millicell EZ SLIDE 8-well glass slides (Merck), respectively. The stable cell lines were induced at 80% confluence by adding a final concentration of 2 μg ml^-1^ doxycycline hydrochloride. The protein overproduction was carried out for 24 h. For permeabilization and staining, cells were washed with PBS twice and fixed in 4% PFA for 15 minutes at room temperature, followed by washing with PBS twice, and permeabilized in PBST (PBS with 0.5% triton-X) for 15 minutes at room temperature. For immunofluorescent staining, the HEK293 cells were subsequently washed with PBS and blocked in 5% normal goat serum for one hour before staining with FLVCR1 and FLVCR2 polyclonal antibodies at 1:250 dilutions for 1 h and then with alexa Fluor 555 as secondary antibody at 1:500 dilutions for 1 h. The cells were counter-stained with DAPI and imaged with laser confocal microscope (Zeiss LSM710). The overproduction stable cells were treated with the same protocol but stained with Monoclonal ANTI-FLAG® M2-FITC antibody against their FLAG-tags, and MitoTracker™ Red CMXRos for mitochondria localization. The cells were mounted using ProLong™ Diamond mounting medium with DAPI and imaged with laser confocal microscope (Confocal Microscope Leica STELLARIS 5).

### Structure-guided mutagenesis

To generate the mutant plasmids for *FLVCR1* and *FLVCR2*, an overlapping PCR approach was used. The mutated cDNA of *FLVCR1* or *FLVCR2* was cloned into pcDNA3.1 for overexpression. The mutations were validated by Sanger sequencing. To test the transport activity of these mutants, the mutant plasmid was either co-transfected with *CHKA* for choline transport assay or *ETNK-1* for ethanolamine transport assay. After 24 hours of post-transfections, cells were washed with DMEM and incubated with DMEM containing 20 μM [^3^H] choline or 2.5 μM [^14^C] ethanolamine for *FLVCR1* mutants and 100 μM [^3^H] choline or 2.5 μM [^14^C] ethanolamine for *FLVCR2* mutants. The assays were stopped after 1 h of incubation at 37 °C. Radioactive signal of each mutant was normalized to the total protein levels.

### Choline export assay

To examine the export function, *FLVCR1* and *FLVCR2* plasmids were expressed in HEK293 cells without co-transfection with *CHKA* or *ETNK-1*. The cells were then incubated with 200 μM [^3^H] choline or 100 μM [^14^C] ethanolamine for 2 hours to prepack the cells with the ligand. Subsequently, the cells were washed to remove the ligands left over in the medium and incubated with choline/ethanolamine-free medium for 1 h at 37 °C for the release of pre-packed ligand. The cells were washed and collected for quantification of radioactive signals. Samples after 2 hours of incubation with the radioactive ligand were collected to determine the levels of radioactive levels before the release and used for control.

### Hemin import assay

For hemin import activity measurement, HEK293 cells were transfected with *FLVCR1* or *FLVCR2*. After 24 hours of transfection, cells were incubated with 2.5 μM [^3^H] hemin in DMEM containing 10% FBS for 1 h. The cells were then washed and collected for quantification of radioactive signals. For the transport assays where hemin was co-incubated with radioactive ligands, HEK293 cells with overexpression of *FLVCR2* were incubated with 50 μM hemin and 10 μM [^3^H] choline or 20 μM hemin and 2.5 μM [^14^C] ethanolamine for 1 hour. Radioactive signals from the cells were quantified.

### Deglycosylation assay

PNGase assay was performed on FLVCR1 and FLVCR2 proteins purified from HEK293 cells were treated with PNGase (NEB). *FLVCR1* and *FLVCR2* expressed in Expi293T GnTi-cells were used for control. Treated protein samples were then subjected to Western blot analysis.

### Metabolomic analysis

Adult livers (aged 3-6 months old) from control (*FLVCR1*^f/f^) and conditional *FLVCR1* knockout (*FLVCR1*^f/f^ Mx1-Cre) mice were used for metabolomic analysis. Briefly, the mice were perfused with PBS to remove blood before organ collection. Liver samples were snap-frozen before shipped for metabolomics by Metabolon. The levels of metabolites were expressed as relative amount.

### Production and purification of the human FLVCR1 and FLVCR2

For protein production, the Flp-In T-REx293-FLVCR1 and Flp-In T-REx293-FLVCR2 cell lines were cultured in roller bottles (Greiner Bio-One) in growth media containing 100 μg ml^-1^ hygromycin B for 14 d under the above-mentioned conditions. Gene expression was induced at 100% confluence by adding a final concentration of 2 μg ml^-1^ doxycycline hydrochloride. After 72 h, cells were harvested with Accutase solution (Sigma-Aldrich) and stored at −80 °C until further use. Harvested cells were suspended in cold lysis buffer containing 25 mM Tris pH 7.4, 150 mM NaCl, and 0.1 g ml^-1^ SigmaFast ethylenediaminetetraacetic acid (EDTA)-free protease inhibitor (Sigma-Aldrich) and disrupted by stirring under high-pressure nitrogen atmosphere (750 MPa) for 45 min at 4 °C in a cell-disruption vessel (Parr Instrument). The cell lysate was centrifuged at 8,000*g* at 4 °C for 15 min. Subsequently, the low-velocity supernatant was centrifuged at 220,000*g* at 4 °C for 60 min. Pelleted membranes were resuspended and stored in a storage buffer containing 25 mM Tris pH 7.4, 150 mM NaCl, 10% glycerol (v/v), and 0.1 g ml^-1^ SigmaFast EDTA-free protease inhibitor (Sigma-Aldrich).

All purification steps of both FLVCRs were performed at 4 °C. Isolated membranes were solubilized with 1% (w/v) lauryl maltose neopentyl glycol (LMNG; GLYCON Biochemicals) with gentle stirring for 1 h. The insoluble membrane fraction was removed via ultracentrifugation at 220,000*g* for 1 h. Subsequently, the supernatant was incubated with ANTI-FLAG® M2 Affinity Gel resin (Merck) for 1 h. The resin was preequilibrated with a buffer containing 50 mM Tris pH 7.4, 150 mM NaCl, and 0.02% LMNG (w/v). The washing step was performed using 20 column volumes (CVs) of wash buffer [50 mM Tris pH 7.4, 150 mM NaCl, 5% (v/v) glycerol, and 0.02% LMNG). The protein was eluted from the M2 resin with 10 CVs of the same buffer supplemented with 4 mM FLAG® Peptide (Merck). The eluted sample was concentrated and subjected to a Superdex 200 Increase 10/300 column (GE Healthcare) equilibrated with size exclusion chromatography (SEC) buffer [50 mM Tris pH 7.4, 150 mM NaCl, and 0.001% (w/v) LMNG]. Peak fractions were pooled, concentrated to 1.5 mg ml^-1^ using an Amicon 50-kDa cut-off concentrator (Merck Millipore), and stored for further analysis.

### Immunoblotting

Affinity-purified proteins were subjected to SDS-PAGE and immunoblotting. FLAG-tagged FLVCR1 and FLVCR2 were detected using anti-FLAG (F3165, Sigma-Aldrich) antibodies at 1:1,000 dilution. Anti-mouse IgG antibody conjugated with alkaline phosphatase (A9316, Sigma-Aldrich) was used as secondary antibody at 1:5,000 dilution. Native FLVCR1 and FLVCR2 proteins were detected by polyclonal FLVCR1 and FLVCR2 antibodies raised in-house at 1:1000 dilution. GAPDH antibody was purchased from Santa Cruz.

### Tryptophan fluorescence measurement

Tryptophan fluorescence measurements were carried out using Prometheus Panta (NanoTemper Technologies). Purified protein samples were diluted with dilution buffer containing 50 mM HEPES pH 7.4, 150 mM NaCl, and 0.001% (w/v) LMNG to 1 μM. Buffers with different concentrations of choline or betaine were prepared by serial dilutions of dilution buffer containing 4 mM of the compounds. The protein samples were mixed with an equal volume of dilution buffer or the compound-containing buffer with a final protein concentration of 0.5 μM and then incubated at room temperature for 15 min. A volume of 10 µl mixed solution was used per Prometheus high sensitivity capillary (NanoTemper Technologies). Recorded *F*_350_/*F*_330_ was analyzed by using Python libraries including pandas, numpy, scipy and seaborn in Visual Studio Code (Microsoft). Three technical replicates were recorded for data analysis.

### Cryo-EM sample preparation

In order to collect cryo-EM data of FLVCR1 and FLVCR2 in different sample conditions, different combinations of FLVCR proteins and putative substrate molecules were prepared. For both as-isolated samples of FLVCRs, the protein concentration was adjusted to approximately 1.5 mg ml^−1^ and subjected to plunge freezing. For samples supplemented with heme, heme loading was performed prior to the SEC during protein purification with a 10-fold molar excess to the protein concentration. Peak fractions were pooled and concentrated to 1.5 mg ml^−1^ before sample vitrification. For samples supplemented with choline, purified proteins were adjusted to 1.5 mg ml^−1^ and choline was added at a final concentration of 1 mM. The samples were incubated for 10 minutes at room temperature before plunge freezing. Identical plunge freezing conditions were applied for all samples: Quantifoil R1.2/1.3 copper grids (mesh 300) were washed in chloroform and subsequently glow-discharged with a PELCO easiGlow device at 15 mA for 90 seconds. A volume of 4 µl sample was applied to a grid and blotting was performed for 4 seconds at 4 °C, 100% humidity with nominal blot force 20 immediately before freezing in liquid ethane, using a Vitrobot Mark IV device (Thermo Scientific).

### Cryo-EM image recording

For each cryo-EM sample, a dataset was recorded in Energy-Filtered Transmission Electron Microscopy (EF-TEM) mode using either a Titan Krios G3i or a Krios G4 microscope (Thermo Scientific), both operated at 300 kV. Electron-optical alignments were adjusted with EPU 3.0 - 3.4 (Thermo Scientific). Images were recorded using automation strategies of EPU 3.0 - 3.4 in electron counting mode with either a Gatan K3 (installed on Krios G3i) or a Falcon4 (installed on Krios G4) direct electron detector. For Gatan K3 detector, a nominal magnification of 105,000, corresponding to a calibrated pixel size of 0.837 Å was applied, and dose fractionated movies (80 frames) were recorded at an electron flux of approximately 15 e^-^ x pixel^-1^ x s^-1^ for 4 s, corresponding to a total dose of ∼80 e^-^/A^2^. For Falcon4 detector, a nominal magnification of 215,000, corresponding to a calibrated pixel size 0.573 Å was applied, dose fractionated movies were recorded in electron-event representation (EER) format at an electron flux of approximately 4 e^-^ x pixel^-1^ x s^-1^ for 5 s, corresponding to a total dose of ∼50 e^-^/A^2^. Images were recorded between −1.1 and −2.0 µm nominal defocus. Data collection quality was monitored through EPU v. 3.0-3.4 and CryoSparc Live (versions 3.0 and 4.0)^38^.

### Cryo-EM image processing

For each acquired dataset, the same cryo-EM image processing approach was applied: MotionCor2 was used to correct for beam-induced motion and to generate dose-weighted images^39^. Gctf was used to determine the contrast transfer function (CTF) parameters and perform correction steps^40^. Images with estimated poor resolution (>4 Å) and severe astigmatism (>400 Å) were removed at this step. Particles were picked by TOPAZ and used for all further processing steps^41^. 2D classification, initial model generation, 3D classification, CTF refinement, Bayesian polishing, 3D sorting, and final map reconstructions were performed using RELION (versions 3.1 and 4.0) or cryoSPARC (versions 3.0 and 4.0)^38,42,43^. Fourier shell correlation (FSC) curves and local-resolution estimation were generated in RELION or cryoSPARC for individual final maps. A schematic overview of our processing workflow, and a summary of map qualities are shown in Supplementary Figs. 2, 3 and 9, and Supplementary Table 4.

### Model building and geometry refinement

The first atomic model of FLVCR1 and FLVCR2 were built into the respective EM density maps of the as-isolated state in Coot (version 0.8) or ISOLDE within ChimeraX (version 1.5 and 1.6)^44–46^, using the AlphaFold predicted structures as initial templates^47^. After manual backbone tracing and docking of side chains, real-space refinement in Phenix was performed (version 1.18)^48^. Refinement results were manually inspected and corrected if required. This model was used as a template to build all subsequent atomic models. The finalized models were validated by MolProbity implemented in Phenix^49^. Map-to-model cross-validation was performed in Phenix (version 1.18). FSC_0.5_ was used as cut-off to define resolution. A summary of model parameters and the corresponding cryo-EM map statistics is found in Supplementary Table 4. The finalized models were visualized using ChimeraX. The built models of both FLVCR proteins in different states were used as starting structures for MD simulations.

### Molecular dynamics simulations

All simulations were run using GROMACS 2022.4^50^. The protein structures were embedded in a lipid bilayer with 75% POPE and 25% POPG with CHARMM-GUI^51^ and solvated in TIP3P water with 150 mM NaCl. The charmm36m force field^52^ was used with the improved WYF parameter set for cation-pi interactions^53^. Initial systems were minimized for 5000 steepest-descent steps and equilibrated for 250 ps of MD in an NVT ensemble and for 1.625 ns in an NPT ensemble. Position restraints of 4000 and 2000 kJ mol^−1^ nm^−2^ in the backbone and side chain heavy atoms, respectively, were gradually released during equilibration. The z-positions of membrane phosphates, as well as lipid dihedrals, were initially restrained with force constants of 1000 kJ mol^−1^ nm^−2^, which were gradually released during equilibration. The initial time step of 1 fs was increased to 2 fs during NPT equilibration. Long-range electrostatic interactions were treated with particle-mesh Ewald (PME)^54^ with a real-space cut-off of 1.2 nm. Van-der-Waals interactions were cut-off beyond a distance of 1.2 nm. The LINCS algorithm^55^ was used to constrain the bonds involving hydrogen atoms. During equilibration, a constant temperature of 310 K was maintained with the Berendsen thermostat^56^, using a coupling constant of 1 ps. Constant pressure of 1 bar was established with a semi-isotropic Berendsen barostat and a coupling constant of 5 ps. In the production runs, a Nosé–Hoover thermostat^57^ and a Parrinello– Rahman barostat were used^58^.

We used our cryo-EM structures as initial models for simulations of apo and choline-bound inward-facing FLVCR1, apo and choline-bound inward-facing FLVCR2, and apo outward-facing FLVCR2. An initial structure of choline-bound outward-facing FLVCR2 was generated aligning apo outward-facing FLVCR2 to choline-bound inward-facing FLVCR2 and maintaining choline in the cavity. In choline entry simulations, the apo structures were used with 380 mM choline in solution. For simulations of ethanolamine-bound FLVCR1, the choline within the cavity of the cryo-EM structure was replaced by this ligand. Simulations of heme-bound outward- and inward-facing FLVCR2 were prepared by manually placing heme close to the observed binding region in the N-terminus of the apo structures, outside of the lipid bilayer. Additionally, simulations of FLVCR2 in the outward-facing conformation were conducted, with the heme initially positioned in front of the cavity entrance. Three replicas with random initial velocities from the Boltzmann distribution were run for all systems. Choline release simulations were interrupted after choline exit from the cavity, and hence have variable duration. For all other systems, each replica was run for 1 µs.

Alanine substitution mutations were introduced using PyMol^59^ and simulated with identical parameters as those applied in the corresponding wild-type simulations. In FLVCR1, only one mutation was introduced in the cavity residue W125. Mutations of FLVCR2 included the cavity residue W102 and the heme binding residues R82, R83 (simultaneously) and K273A.

Visual Molecular Dynamics (VMD)^60^ and MDAnalysis^61^ were used to visualize and analyze the trajectories, respectively.

### Interior tunnels and cavities

Tunnels and cavities were mapped with MOLE 2.5^62^ with a bottleneck radius of 1.2 Å, bottleneck tolerance 3 Å, origin radius 5 Å, surface radius 10 Å, probe radius 5 Å and an interior threshold of 1.1 Å. We calculated the volume of the cavity using CASTp^63^ with a bottleneck radius of 1.4 Å. Residues 297-320 and 512-516 were removed from the FLVCR1 model to avoid the misattribution of the volume between internal loops to the cavity volume. Analogously, residues 272-296 and 487-502 were not included in the cavity volume calculation of FLVCR2.

### Multiple sequence alignments

Multiple sequence alignments of FLVCR1 and FLVCR2 from *homo sapiens*, *Felis catus*, *Mus musculus*, and *Sus scrofa* were performed using Clustal Omega^64^.

## Data availability

Cryo-EM maps are deposited at the Electron Microscopy Data Bank under accession numbers:<u>EMD-</u> <u>18334</u>, <u>EMD-18335</u>, <u>EMD-18336</u>, <u>EMD-18337</u>, <u>EMD-18338</u>, <u>EMD-18339</u>, <u>EMD-19009</u>, <u>EMD-19018</u>. Atomic models of human FLVCR1 and FLVCR2 have been deposited to the Protein Data Bank under accession numbers: <u>8QCS</u>, <u>8QCT</u>, <u>8QCX</u>, <u>8QCY</u>, <u>8QCZ</u>, <u>8QD0</u>, <u>8R8T</u>. All other data is presented in the main text or supplementary materials. Source data are provided with this paper.

## Acknowledgements

We thank Hartmut Michel for supporting and providing infrastructural resources. We thank the Central Electron Microscopy Facility at MPI of Biophysics for technical support and access to instrumentation.

## Author contributions

T-H.W. purified proteins, performed biochemical assays, prepared grids, collected cryo-EM data, processed cryo-EM data, refined the structure, built the model, co-drafted the manuscript, and prepared figures. K.R., Z.Y., and T.J.Y. L. performed mutagenesis. K.R. performed transport assays and WB for mutants, and export assays. N.C.P.L. and Z.Y. performed dose curve, time course, and glycosylation assay, and immunostaining. A.C.C. performed MD simulations, analyzed data, co-drafted the manuscript, and prepared figures. W.J. performed cell productions, optimized purification conditions, and purified proteins. A.B. performed assays and analyzed data. R.T.D and J.L.A designed and performed animal model experiments. S.W. calibrated and aligned the microscope. G.G. performed IF assays and protein purification. S.L.S performed protein purification. G.H., D.W., L.N.N., and S.S. supervised the project. D.W. implemented cell production and protein purification, prepared grids, performed initial cryo-EM screening experiments, collected cryo-EM data, analyzed data, drafted the manuscript, generated figures, and funded the project. L.N.N. designed functional assays, interpreted, analyzed data, funded the project, and co-drafted the manuscript. D.W. and S.S. initiated the project. S.S. designed research, evaluated data, funded the project, drafted the manuscript, and generated figures.

---

**Extended Data Fig. 1:**
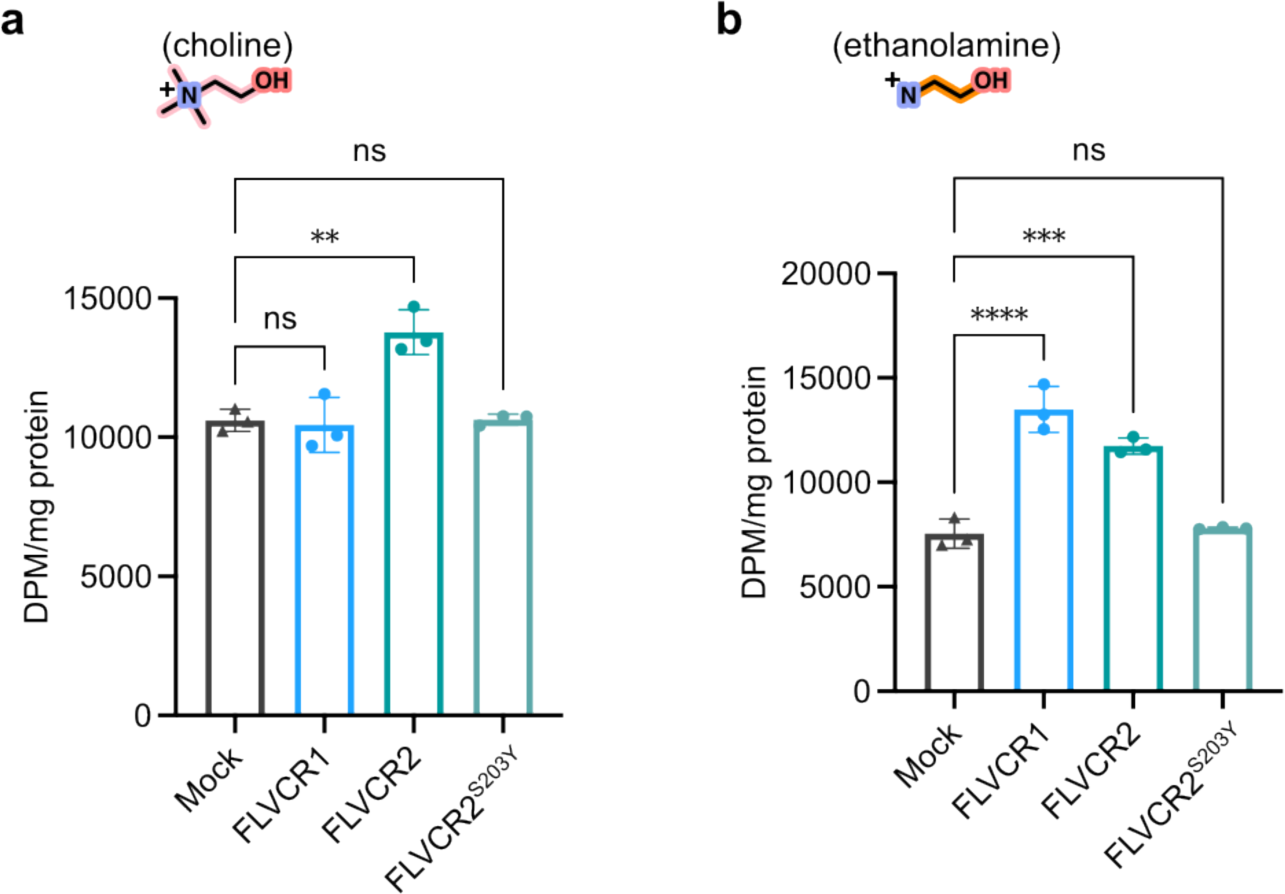
Transport activity of FLVCRs for choline or ethanolamine without overexpression of *CHKA* or *ETNK-1*. Choline (**a**) or ethanolamine (**b**) transport activity of FLVCR1, FLVCR2, and FLVCR2^S203Y^ without simultaneous overexpression of *CHKA* or *ETNK-1*, respectively. 20µM [^3^H] choline or 2.5µM [^14^C] ethanolamine was used, respectively. The inactive mutant of FLVCR2^S203Y^ was used as a control in all experiments. Each experiment was repeated twice and one dataset was shown. Each symbol represents one replicate. Data are expressed as mean ± SD. ****P<0.0001, ***P<0.001, **P<0.01. ns, not significant. One-way ANOVA.

**Extended Data Fig. 2:**
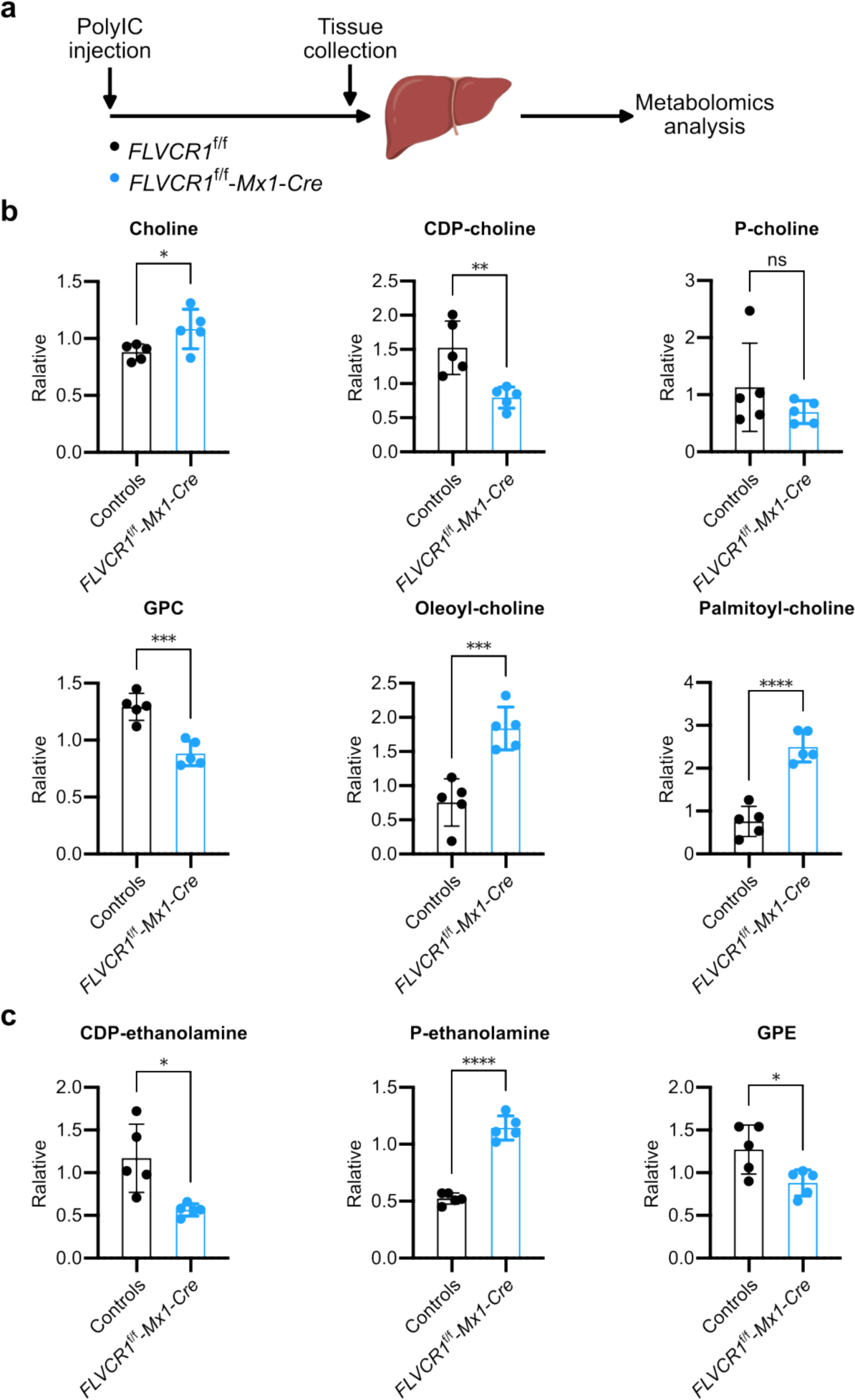
Metabolomic analysis of livers from *FLVCR1*-knockout mice. **a**, Illustration of experimental procedures. *FLVCR1* was deleted by polyIC injection in *FLVCR1*^f/f^-Mx1-Cre (knockout) mice. Liver samples of control (*FLVCR1*^+/+^-Mx1-Cre) and knockout mice were collected at least 4 weeks post-injection for metabolomic analysis. **b,** Levels of choline and choline metabolites from control and knockout mice. **c**, Levels of ethanolamine metabolites from control and knockout mice. Each data point represents one mouse. Data are expressed as mean ± SD. ****P<0.0001, ***P<0.001, **P<0.01, *P<0.05. ns, not significant. *t*-test.

**Extended Data Fig. 3:**
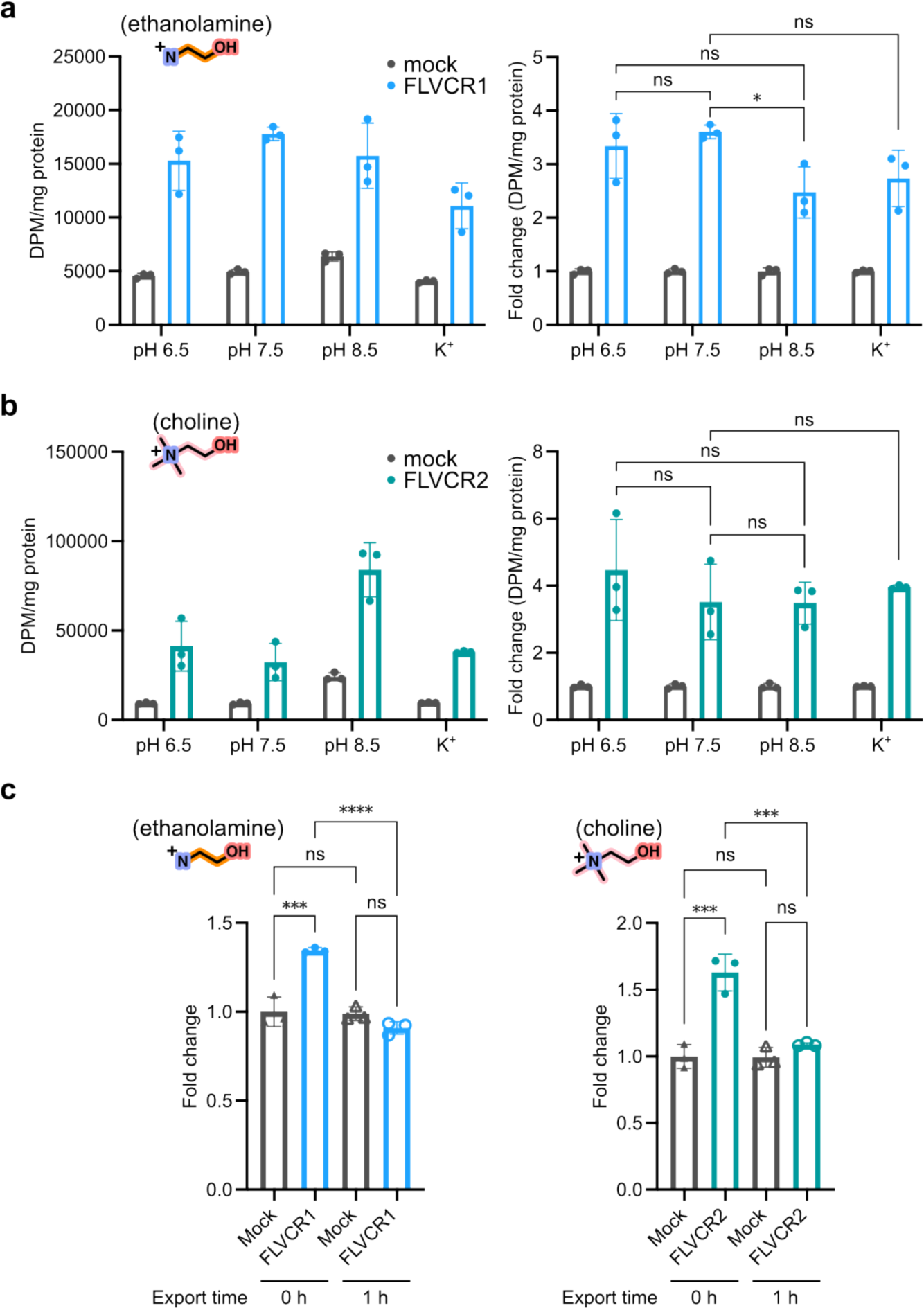
Transport properties of FLVCR1 and FLVCR2. **a,** Ethanolamine transport activity of FLVCR1 at indicated pH values, and in sodium-free buffer (K^+^). **b**, Choline transport activity of FLVCR2 at indicated pH values, and in sodium-free buffer (K^+^). The right panels show normalized data with reference to the respective mocks. In these experiments, 2.5 µM [^14^C] ethanolamine was used for FLVCR1 or 20 µM [^3^H] choline for FLVCR2, respectively. The cells were co-expressed with *ETNK-1* or *CHKA* and incubated with the ligands at 37 °C for 15 mins. **c**, Export assays of FLVCR1 with ethanolamine (left) and FLVCR2 with choline (right). In these assays, 100 µM [^14^C] ethanolamine or 200 µM [^3^H] choline was incubated with *FLVCR1* or *FLVCR2* overexpression cells, respectively. After 2 hours of incubation, the buffer was washed out. Intracellular [^3^H] choline or [^14^C] ethanolamine from the cells was allowed to release into choline/ethanolamine-free medium for 1 hour. The radioactive signal in the cells was normalized to the total protein and expressed as fold change to mock. Each symbol represents one replicate. Data are expressed as mean ± SD. ****P<0.0001, ***P<0.001, *P<0.05. ns, not significant. Two-way ANOVA for (**a**) and (**b**), One-way ANOVA for (**c**).

**Extended Data Fig. 4:**
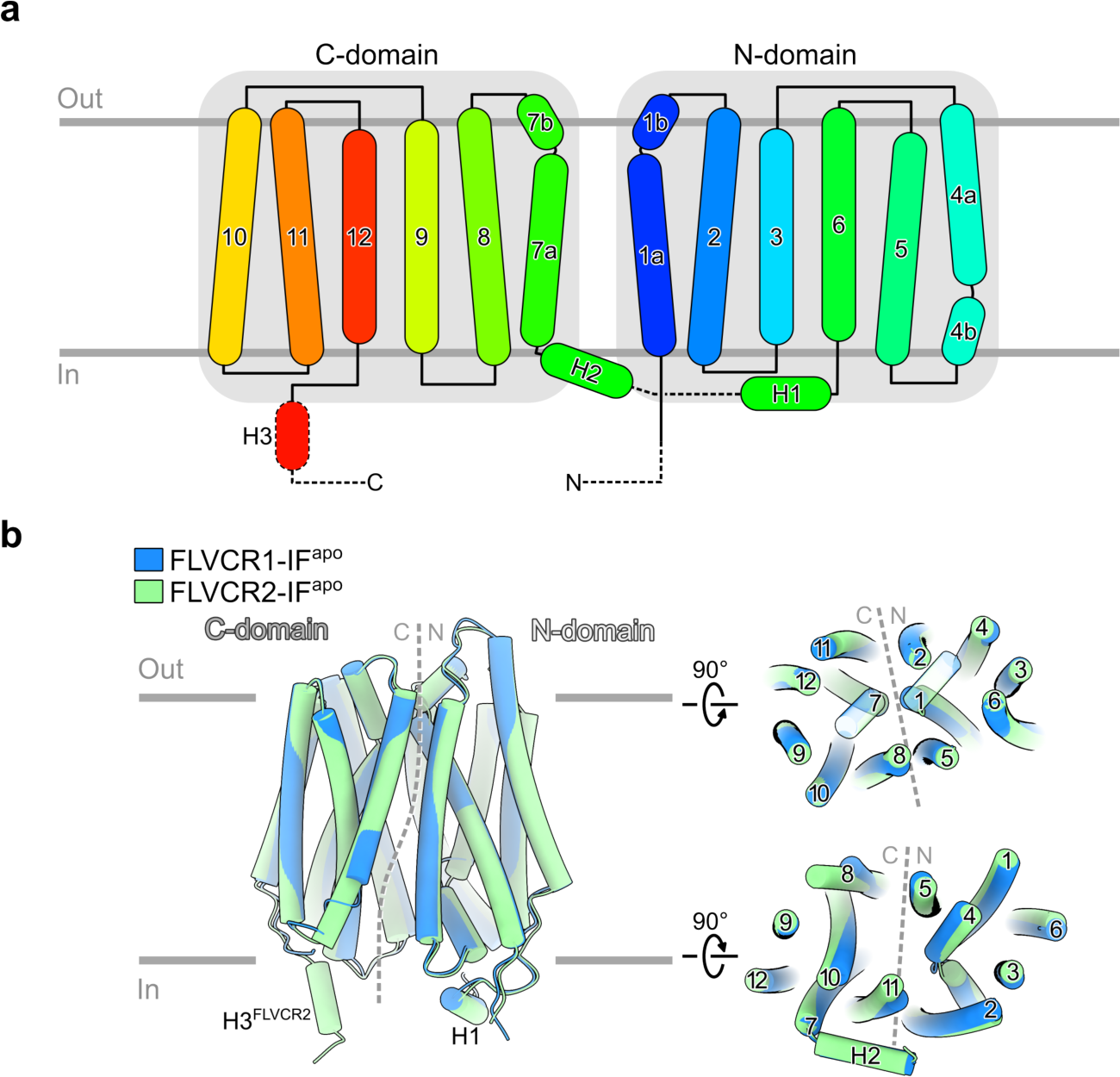
Architecture and structural comparison between FLVCR1 and FLVCR2. **a**, Schematic diagram of FLVCR family showing the topology of the secondary structure. Motifs that are not observed in both cryo-EM structures of FLVCR1 and FLVCR2 are shown as dashed lines. **b**, Structural comparison of FLVCR1-IF^apo^ and FLVCR2-IF^apo^ in tube representation, viewed from the lipid bilayer (left), extracellular side (top-right), and intracellular side (bottom-right).

**Extended Data Fig. 5:**
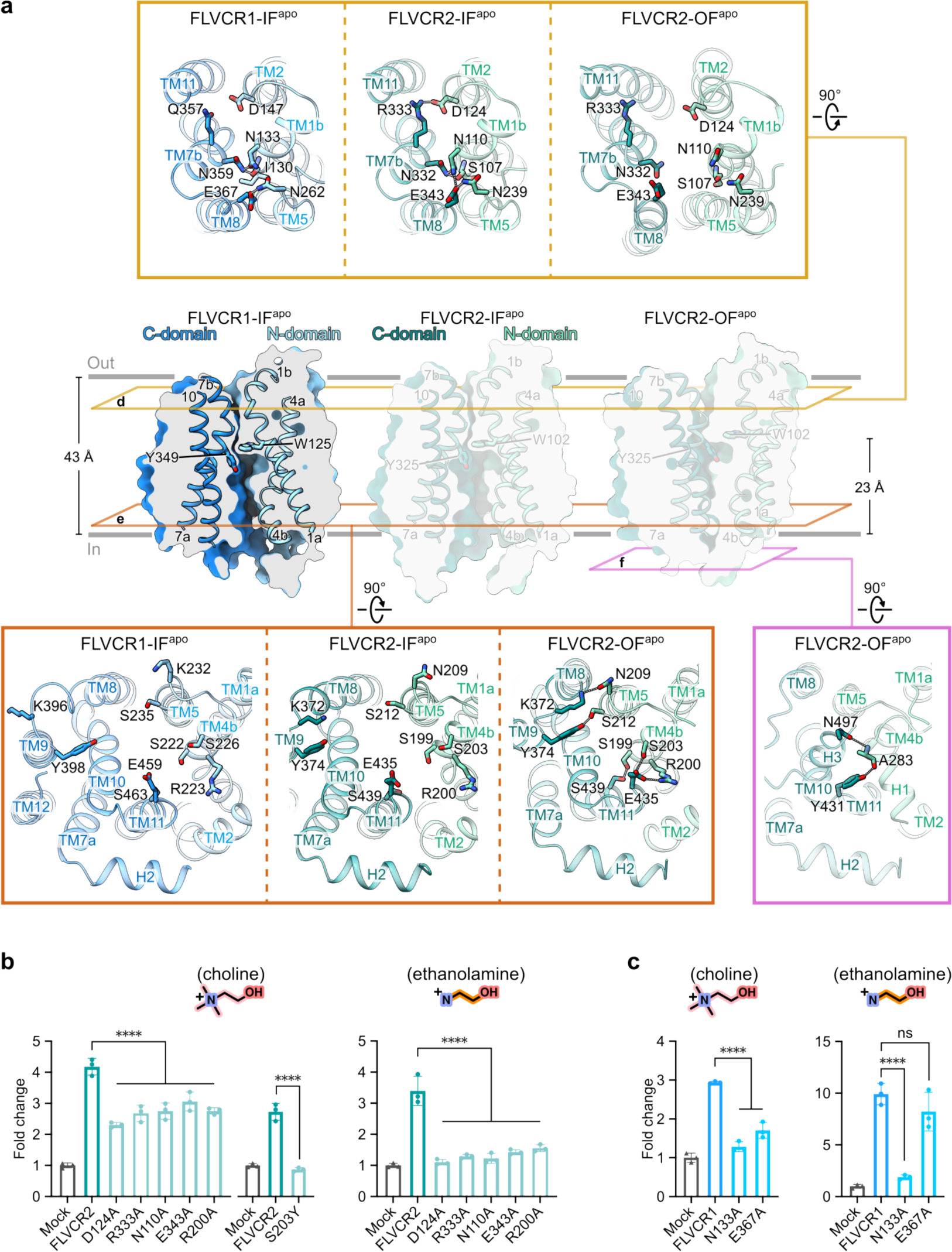
Residues for intra- and extracellular gating in FLVCR1 and FLVCR2. **a**, Cut-away view of FLVCR1-IF^apo^ in surface representation showing the cytoplasmic cavity. Two central aromatic residues are shown as sticks. The structures of FLVCR2-IF^apo^ and FLVCR2-OF^apo^ are shown for comparison. Cross-sections of their inter-domain interactions are shown from the extracellular side (top), or from the intracellular side (bottom-left). Residues in FLVCR1 corresponding to the inter-domain interaction residues in FLVCR2 are shown. The bottom-right panel shows the inter-domain interactions between H1 and H3 in FLVCR2-OF^apo^ viewed from the intracellular side. Residues participating in the inter-domain interactions are shown as sticks; hydrogen bonds and salt bridges are labelled with dashed lines. **b**, Transport assay of FLVCR2 mutants for choline (left) and ethanolamine (right). *CHKA* or *ETNK-1* was co-expressed with wild-type *FLVCR2* and mutant plasmids. In these assays, 100 µM [^3^H] choline or 2.5 µM [^14^C] ethanolamine was used, respectively. **c**, Transport assay of FLVCR1 mutants for choline (left) and ethanolamine (right). *CHKA* or *ETNK-1* was co-expressed with wild-type *FLVCR1* and mutant plasmids, and 20 µM [^3^H] choline or 2.5 µM [^14^C] ethanolamine was used, respectively. Each symbol represents one replicate. Data shown are mean ± SD. ****P<0.0001. ns, not significant. One-way ANOVA.

**Extended Data Fig. 6:**
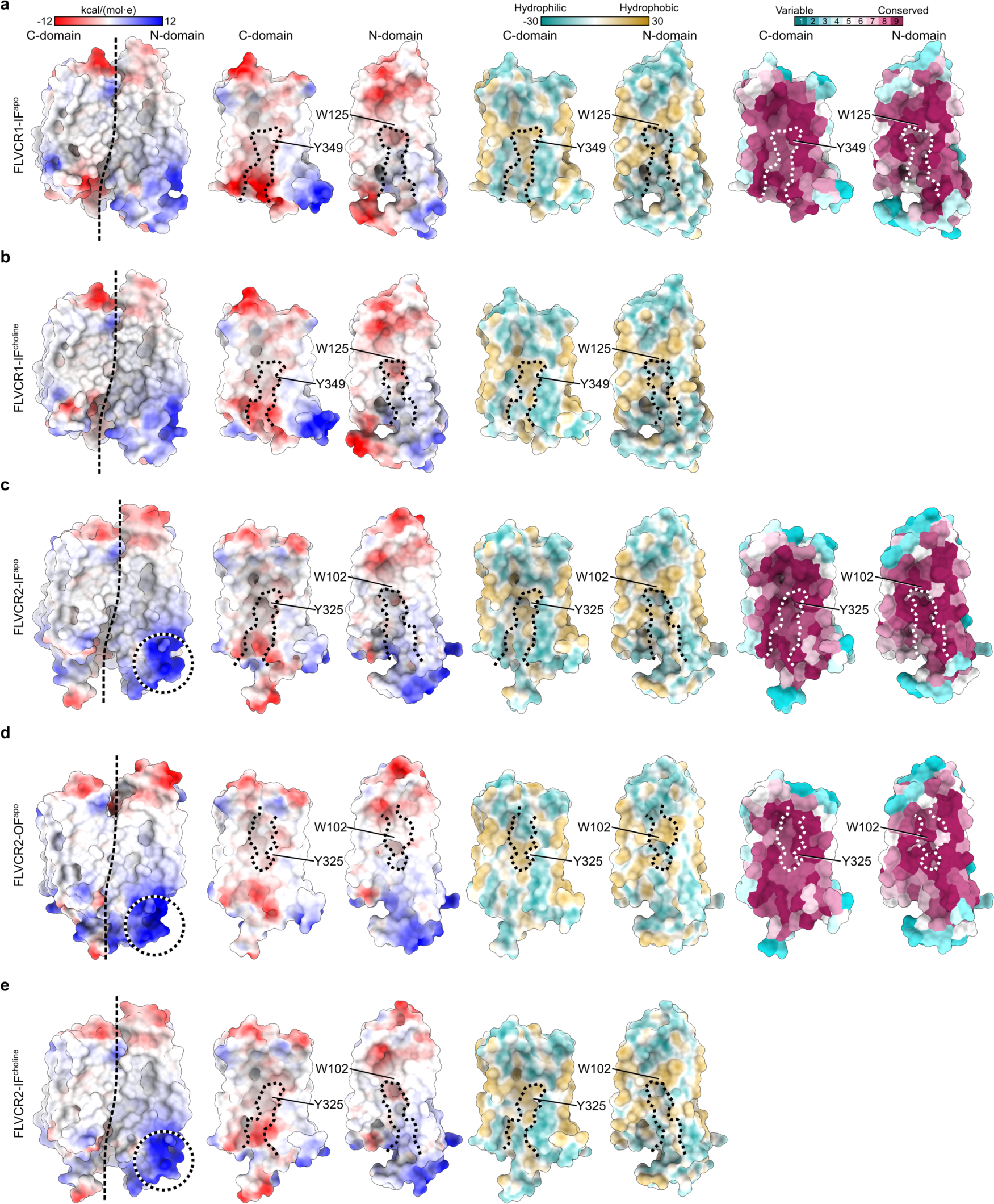
Physicochemical properties of FLVCR structures and conservation analyses. Surface charge, hydrophobicity, and conservation analyses of FLVCR1-IF^apo^ (**a**), FLVCR1-IF^choline^ (**b**), FLVCR2-IF^apo^ (**c**), FLVCR2-OF^apo^ (**d**), and FLVCR2-IF^choline^ (**e**). From left to right: Surface viewed from the lipid bilayer and both C and N domains viewed from the domain interface coloured by electrostatic potential, the domain interface coloured by hydrophobicity, and the domain interface coloured by sequence conservation. The cavity in the different states of both FLVCRs is outlined with dashed lines. Two important central pocket residues (W125 and Y349 in FLVCR1; W102 and Y325 in FLVCR2) are indicated. The FLVCR2 binding site for heme is marked by black-and-white circles.

**Extended Data Fig. 7:**
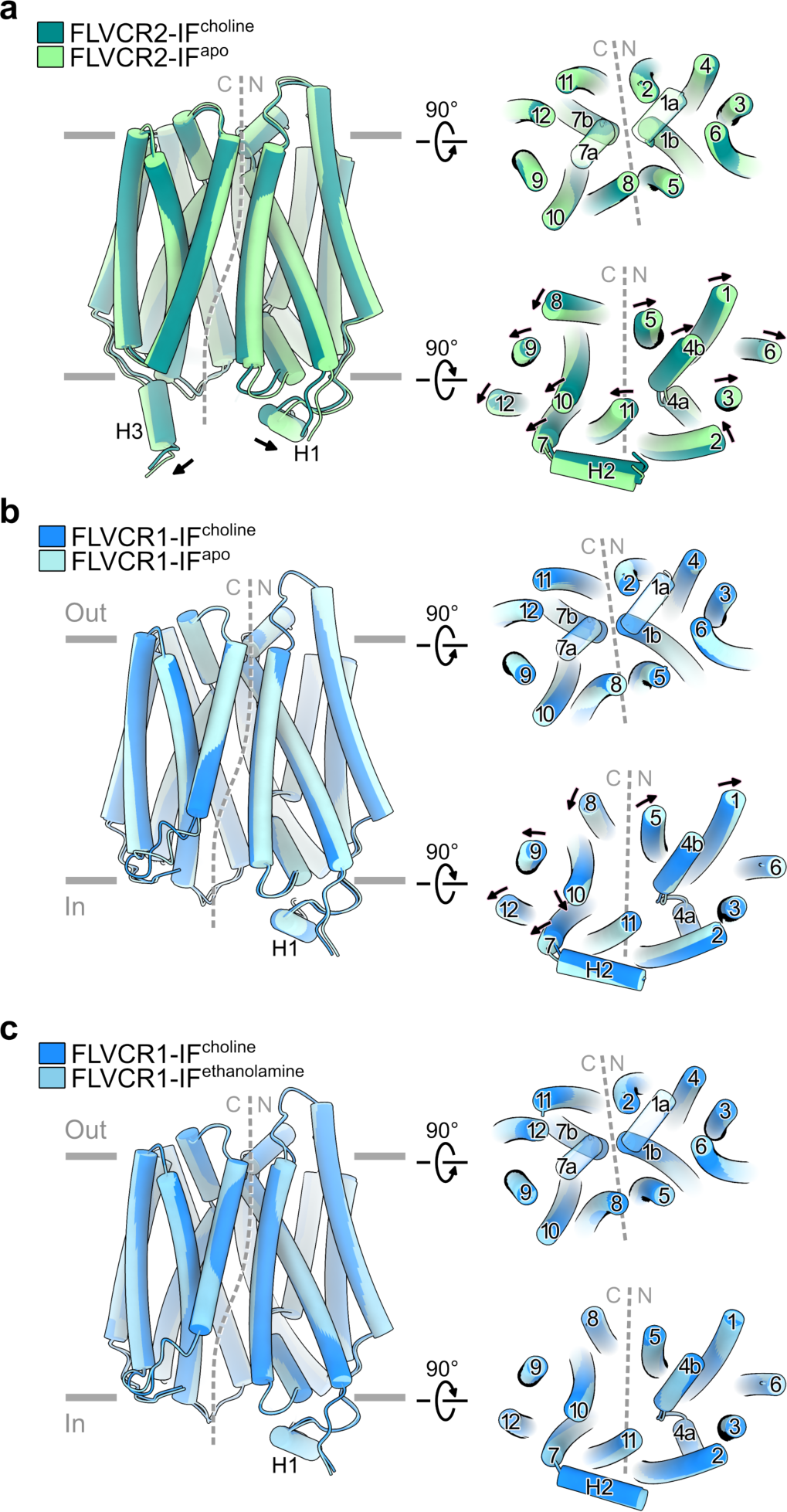
Conformational changes of substrate-bound FLVCRs. Structural comparison of FLVCR2-IF^choline^ and FLVCR2-IF^apo^ (**a**), FLVCR1-IF^choline^ and FLVCR1-IF^apo^ (**b**), as well as FLVCR1-IF^choline^ and FLVCR1-IF^ethanolamine^ (**c**), viewed from the lipid bilayer (left), extracellular side (top-right), and intracellular side (bottom-right).

**Extended Data Fig. 8:**
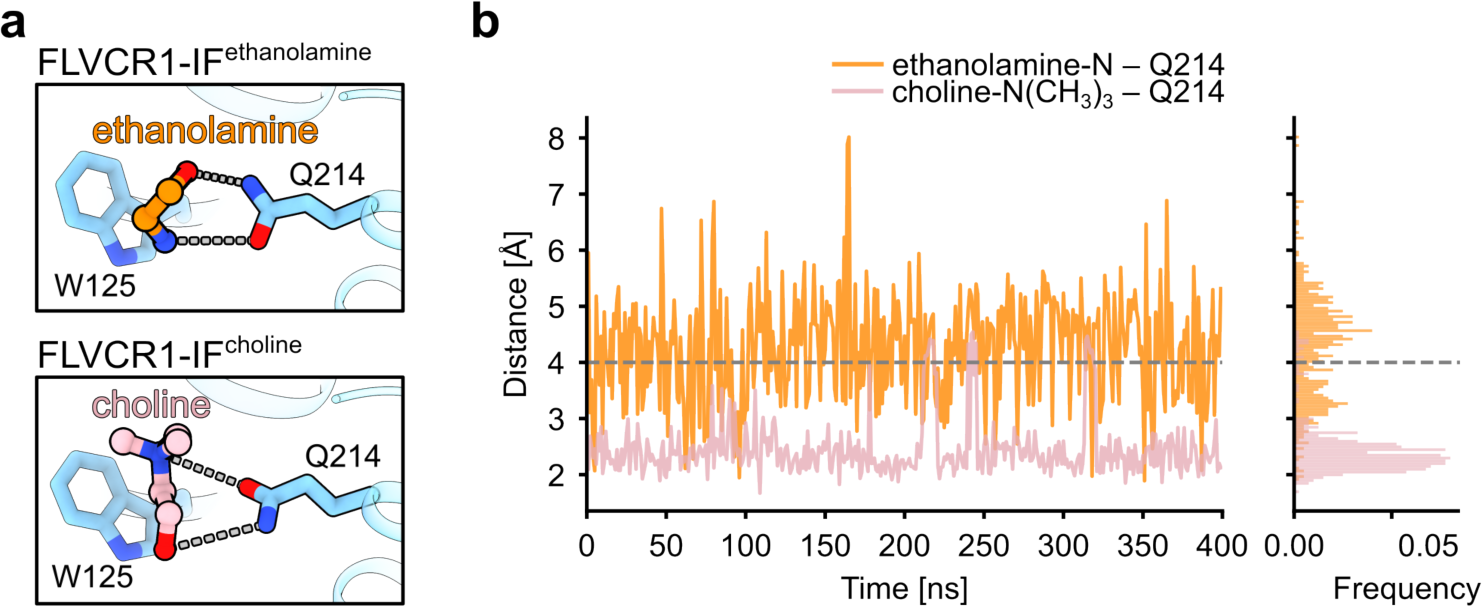
Interaction chemistries and dynamics between Q214F^LVCR1^ and respective ligands. **a**, Structural comparison of the binding site of ethanolamine- and choline-bound FLVCR1. **b**, Time-resolved distance plots between Q214 and the primary amine of ethanolamine (orange) or the tertiary amine of choline (pink) obtained from MD simulation runs. The right panel shows the accumulative frequency of their distances during the simulations.

**Extended Data Fig. 9:**
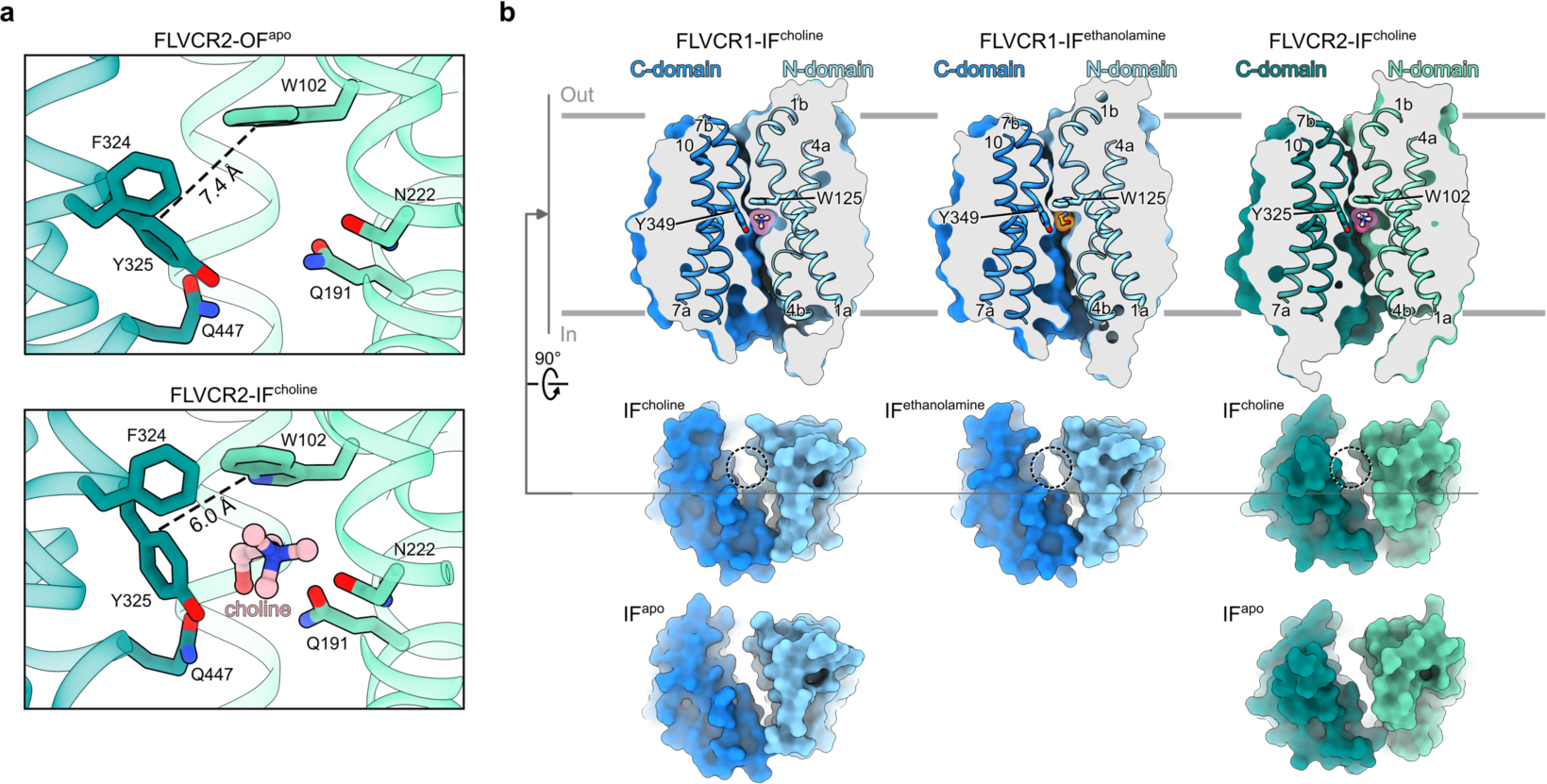
Translocation pathway of choline in FLVCRs. **a**, Choline-binding sites of FLVCR2-OF^apo^ (top) and FLVCR2-IF^choline^ (bottom) with the distance between W102 and Y325 shown as dashed lines. **b**, Cut-away views of FLVCR1-IF^choline^ (top-left), FLVCR1-IF^ethanolamine^ (top-middle) and FLVCR2-IF^choline^ (top-right) showing the inward-facing cavity. Two central aromatic residues are shown as sticks. The surfaces shown below are the respective models viewed from the intracellular side. The surfaces of IF^apo^ of FLVCR1 and FLVCR2 are also shown for comparison. The dashed circles indicate the peripheral channel in the ligand-bound state.

