## Supplementary information for "Structural and mechanistic insights into human choline and ethanolamine transport"

**Supplementary Table 1 – Nucleotide sequences of human FLVCR variants used in this study.**

| Variant | Nucleotide sequence |
|---|---|
| FLVCR1 (wild-type) | gacggatcgggagatctcccgatcccctatggtgcactctcagtacaatctgctctgatgccgcatagttaagccagtatctgctccctgcttgtgtgttggaggtcg<br>ctgagtagtgcgcgagcaaaatttaagctacaacaaggcaaggcttgaccgacaattgcatgaagaatctgcttagggttaggcgttttgcgctgcttcgcgatgt<br>acgggccagatatacgcgttgacattgattattgactagttattaatagtaatcaattacggggtcattagttcatagcccatatatggagttccgcgttacataact<br>tacggtaaatggcccgcctggctgaccgcccaacgacccccgcccattgacgtcaataatgacgtatgttcccatagtaacgccaatagggactttccattgacgt<br>caatgggtggagtatttacggtaaactgcccacttggcagtacatcaagtgtatcatatgccaagtacgccccctattgacgtcaatgacggtaaatggcccgcct<br>ggcattatgcccagtacatgaccttatgggactttcctacttggcagtacatctacgtattagtcatcgctattaccatggtgatgcggttttggcagtacatcaatg<br>ggcgtggatagcggtttgactcacggggatttccaagtctccaccccattgacgtcaatgggagtttgttttggcaccaaaatcaacgggactttccaaaatgtcgt<br>aacaactccgccccattgacgcaaatgggcggtaggcgtgtacggtgggaggtctatataagcagagctctccctatcagtgatagagatctccctatcagtgat<br>agagatcgtcgacgagctcgtttagtgaaccgtcagatcgcctggagacgccatccacgctgttttgacctccatagaagacaccgggaccgatccagcctccgg<br>actctagcgtttaaacttaagcttgccaccatggcgcggccagacgatgaggagggggcggcggtggcgcccggacacccgctcgcgaaaggatacctcccgtt<br>gccgaggggcgcgcccgttgggaaggagagcgtggagctgcagaacgggcccaaagcgggcaccttcccggtgaatggggcccccgggacagcctcgctgc<br>cgcctcgggagttctgggcgggcctcagactccactggccccagaagaggagacccaggcccggctgctgcctgcgggcgcgggagctgagaccccgggggcc<br>gagagcagcccgctgccccttacggcgctctccccgcggcgcttcgtggtgctcctgatcttcagcctgtactcgctggtcaacgcctttcagtggatccagtacag<br>catcattagcaacgtcttcgagggcttctacggtgtcaccttgctgcacatcgactggctgtccatggtgtacatgctggcctacgtgcccctcatcttcccggccac<br>ctggctgctggacaccagaggcctgcggctcaccgccctgctgggctccggcctcaactgcctgggtgcctggatcaagtgcggcagtgtgcagcagcatctcttc<br>tgggtcaccatgttgggccagtgcttgtgctcggtggcccaggtgttcatcctgggcttgccctcccgcatcgcctcagtgtggtttgggcccaaagaggtgtccac<br>agcttgtgccaccgccgtgctgggcaatcagcttggaactgcagttggctttttgctaccaccagttttagtacccaacacacagaatgacacaaatctcctggctt<br>gtaatatcagcaccatgttttatggaacatcagctgttgccacacttttatttattttaacagcaattgccttcaaagaaaaacctcggtatccaccaagtcaggctc<br>aagcagctcttcaagacagtcccccctgaagagtactcctataagaaatcaataagaaacctgtttaaaaacattccttttgtccttctgttgatcacttatggtatca<br>tgactggtgccttttattcagtctcaacgttattaaatcaaatgatattgacatattatgagggagaagaagtcaatgctggaaggattgggctaacgctagtagta<br>gctggaatggtgggctctattctttgtggcttatggctggattatactaaaacatacaaacagactactctgatagtttatattttgtcttttattggaatggttatcttt<br>actttcacattggaccttagatatattatcatcgtgtttgttactggaggggtgcttggcttcttcatgactggttacctccctttgggttttgaatttgctgttgaaatc<br>acttaccctgaatctgaaggtacttcatctggtcttcttaatgcttctgcacagatatttggaattttgttcacattggctcaaggaaagctcacatcagactatggtc<br>ctaaggcagggaacatttttctctgtgtctggatgtttataggcatcatattaacagcattaatcaagtctgatctgcgaagacacaacataaatataggaattaca<br>aatgttgatgttaaagctataccagctgacagtcccacagaccaagaaccaaaaacggttatgttgtccaagcagtcagaatcagcaattgattacaaggatga<br>cgacgataagtgagcggccgctcgagtctagagggcccgtttaaacccgctgatcagcctcgactgtgccttctagttgccagccatctgttgtttgcccctcccc<br>gtgccttccttgaccctggaaggtgccactcccactgtcctttcctaataaaatgaggaaattgcatcgcattgtctgagtaggtgtcattctattctggggggtggg<br>gtggggcaggacagcaagggggaggattgggaagacaatagcaggcatgctggggatgcggtgggctctatggcttctgaggcggaaagaaccagctggggc<br>tctagggggtatccccacgcgccctgtagcggcgcattaagcgcggcgggtgtggtggttacgcgcagcgtgaccgctacacttgccagcgccctagcgcccgct<br>cctttcgctttcttcccttcctttctcgccacgttcgccggctttccccgtcaagctctaaatcgggggctccctttagggttccgatttagtgctttacggcacctcgac<br>cccaaaaaacttgattagggtgatggttcacgtacctagaagttcctattccgaagttcctattctctagaaagtataggaacttccttggccaaaaagcctgaact<br>caccgcgacgtctgtcgagaagtttctgatcgaaaagttcgacagcgtctccgacctgatgcagctctcggagggcgaagaatctcgtgctttcagcttcgatgta<br>ggagggcgtggatatgtcctgcgggtaaatagctgcgccgatggtttctacaaagatcgttatgtttatcggcactttgcatcggccgcgctcccgattccggaagt<br>gcttgacattggggaattcagcgagagcctgacctattgcatctcccgccgtgcacagggtgtcacgttgcaagacctgcctgaaaccgaactgcccgctgttctg<br>cagccggtcgcggaggccatggatgcgatcgctgcggccgatcttagccagacgagcgggttcggcccattcggaccgcaaggaatcggtcaatacactacatg<br>gcgtgatttcatatgcgcgattgctgatccccatgtgtatcactggcaaactgtgatggacgacaccgtcagtgcgtccgtcgcgcaggctctcgatgagctgatgc<br>tttgggccgaggactgccccgaagtccggcacctcgtgcacgcggatttcggctccaacaatgtcctgacggacaatggccgcataacagcggtcattgactgga<br>gcgaggcgatgttcggggattcccaatacgaggtcgccaacatcttcttctggaggccgtggttggcttgtatggagcagcagacgcgctacttcgagcggaggc<br>atccggagcttgcaggatcgccgcggctccgggcgtatatgctccgcattggtcttgaccaactctatcagagcttggttgacggcaatttcgatgatgcagcttgg<br>gcgcagggtcgatgcgacgcaatcgtccgatccggagccgggactgtcgggcgtacacaaatcgcccgcagaagcgcggccgtctggaccgatggctgtgtag<br>aagtactcgccgatagtggaaaccgacgccccagcactcgtccgagggcaaaggaatagcacgtactacgagatttcgattccaccgccgccttctatgaaagg<br>ttgggcttcggaatcgttttccgggacgccggctggatgatcctccagcgcggggatctcatgctggagttcttcgcccaccccaacttgtttattgcagcttataat<br>ggttacaaataaagcaatagcatcacaaatttcacaaataaagcatttttttcactgcattctagttgtggtttgtccaaactcatcaatgtatcttatcatgtctgta<br>taccgtcgacctctagctagagcttggcgtaatcatggtcatagctgtttcctgtgtgaaattgttatccgctcacaattccacacaacatacgagccggaagcata<br>aagtgtaaagcctggggtgcctaatgagtgagctaactcacattaattgcgttgcgctcactgcccgctttccagtcgggaaacctgtcgtgccagctgcattaatg<br>aatcggccaacgcgcggggagaggcggtttgcgtattgggcgctcttccgcttcctcgctcactgactcgctgcgctcggtcgttcggctgcggcgagcggtatca<br>gctcactcaaaggcggtaatacggttatccacagaatcaggggataacgcaggaaagaacatgtgagcaaaaggccagcaaaaggccaggaaccgtaaaaa<br>ggccgcgttgctggcgtttttccataggctccgcccccctgacgagcatcacaaaaatcgacgctcaagtcagaggtggcgaaacccgacaggactataaagat<br>accaggcgtttccccctggaagctccctcgtgcgctctcctgttccgaccctgccgcttaccggatacctgtccgcctttctcccttcgggaagcgtggcgctttctca<br>tagctcacgctgtaggtatctcagttcggtgtaggtcgttcgctccaagctgggctgtgtgcacgaaccccccgttcagcccgaccgctgcgccttatccggtaact<br>atcgtcttgagtccaacccggtaagacacgacttatcgccactggcagcagccactggtaacaggattagcagagcgaggtatgtaggcggtgctacagagttct<br>tgaagtggtggcctaactacggctacactagaagaacagtatttggtatctgcgctctgctgaagccagttaccttcggaaaaagagttggtagctcttgatccgg<br>caaacaaaccaccgctggtagcggtggtttttttgtttgcaagcagcagattacgcgcagaaaaaaaggatctcaagaagatcctttgatcttttctacggggtct<br>gacgctcagtggaacgaaaactcacgttaagggattttggtcatgagattatcaaaaaggatcttcacctagatccttttaaattaaaaatgaagttttaaatcaa<br>tctaaagtatatatgagtaaacttggtctgacagttaccaatgcttaatcagtgaggcacctatctcagcgatctgtctatttcgttcatccatagttgcctgactccc<br>cgtcgtgtagataactacgatacgggagggcttaccatctggccccagtgctgcaatgataccgcgagacccacgctcaccggctccagatttatcagcaataaa<br>ccagccagccggaagggccgagcgcagaagtggtcctgcaactttatccgcctccatccagtctattaattgttgccgggaagctagagtaagtagttcgccagtt<br>aatagtttgcgcaacgttgttgccattgctacaggcatcgtggtgtcacgctcgtcgtttggtatggcttcattcagctccggttcccaacgatcaaggcgagttaca<br>tgatcccccatgttgtgcaaaaaagcggttagctccttcggtcctccgatcgttgtcagaagtaagttggccgcagtgttatcactcatggttatggcagcactgcat<br>aattctcttactgtcatgccatccgtaagatgcttttctgtgactggtgagtactcaaccaagtcattctgagaatagtgtatgcggcgaccgagttgctcttgcccg<br>gcgtcaatacgggataataccgcgccacatagcagaactttaaaagtgctcatcattggaaaacgttcttcggggcgaaaactctcaaggatcttaccgctgttg<br>agatccagttcgatgtaacccactcgtgcacccaactgatcttcagcatcttttactttcaccagcgtttctgggtgagcaaaaacaggaaggcaaaatgccgcaa<br>aaaagggaataagggcgacacggaaatgttgaatactcatactcttcctttttcaatattattgaagcatttatcagggttattgtctcatgagcggatacatatttg<br>aatgtatttagaaaaataaacaaataggggttccgcgcacatttccccgaaaagtgccacctgacgtc |
| FLVCR2 (wild-type) | gacggatcgggagatctcccgatcccctatggtgcactctcagtacaatctgctctgatgccgcatagttaagccagtatctgctccctgcttgtgtgttggaggtcg<br>ctgagtagtgcgcgagcaaaatttaagctacaacaaggcaaggcttgaccgacaattgcatgaagaatctgcttagggttaggcgttttgcgctgcttcgcgatgt<br>acgggccagatatacgcgttgacattgattattgactagttattaatagtaatcaattacggggtcattagttcatagcccatatatggagttccgcgttacataact<br>tacggtaaatggcccgcctggctgaccgcccaacgacccccgcccattgacgtcaataatgacgtatgttcccatagtaacgccaatagggactttccattgacgt<br>caatgggtggagtatttacggtaaactgcccacttggcagtacatcaagtgtatcatatgccaagtacgccccctattgacgtcaatgacggtaaatggcccgcct |

ggcattatgcccagtacatgaccttatgggactttcctacttggcagtacatctacgtattagtcatcgctattaccatggtgatgcggttttggcagtacatcaatg
ggcgtggatagcggtttgactcacggggatttccaagtctccaccccattgacgtcaatgggagtttgttttggcaccaaaatcaacgggactttccaaaatgtcgt
aacaactccgccccattgacgcaaatgggcggtaggcgtgtacggtgggaggtctatataagcagagctctccctatcagtgatagagatctccctatcagtgat
agagatcgtcgacgagctcgtttagtgaaccgtcagatcgcctggagacgccatccacgctgttttgacctccatagaagacaccgggaccgatccagcctccgg
actctagcgtttaaacttaagcttgccaccatggtgaatgaaggtcccaaccaggaagagagcgatgacacccctgtgccggagtccgcactccaagcggaccc
cagcgtctcggtccatcccagcgtctcggtccatcccagcgtctccatcaaccccagcgtctctgtccaccccagcagttcggcccaccccagtgccttagcccaac
ccagtggcttggctcaccccagtagctcgggccctgaggacctcagcgtgatcaaggtgagcaggcgccgttgggccgtggtcctggtgtttagctgctactccat
gtgcaactcctttcagtggatccagtacggctccatcaataacatcttcatgcacttctacggtgtcagtgcctttgccattgactggctgtccatgtgctacatgctg
acttacatccctctgctcctgccagtggcttggctgctggagaagttcggcctgcgcaccattgctctcactggctcggctctcaactgcctgggggcctgggtgaa
gctgggcagcctgaagccgcatctctttccggtcaccgtggtgggccagctcatctgctctgtggcccaggttttcatcctgggcatgccctcccgcatcgcttccgt
ctggttcggggctaatgaggtttcaacagcctgctccgtggctgtctttggcaatcagcttggaattgcgattgggttcttggtccctcctgttttggtacccaacatt
gaagaccgggacgagcttgcctaccacatcagcatcatgttctatataataggaggtgtggccactctcctcctcatccttgtcatcattgtgttcaaggagaaacc
taaatatcccccagcagggcccaatccctgagctatgccttgacctctcctgatgcctcatacttaggttccatcgcccggctcttcaaaaatctcaactttgtgct
gcttgtcatcacctatggtctgaatgctggtgctttttatgccttgtccactcttctgaatcgcatggtgatctggcactacccgggggaagaagtgaatgctggaag
aattggcctgacgatcgtcattgcaggaatgcttggggctgtgatctcaggaatctggctggataggtccaaaacctacaaagagacaaccctggtagtctatatc
atgacactggtgggcatggtggtgtacacgtttaccttgaacctgggacacctgtgggtagtgttcatcactgctggcacaatgggcttctttatgactggctatctc
ccactgggatttgagtttgctgtggagctcacgtacccagaatcagaaggcatctcctccggcctcctcaacatatctgcacaggtatttgggatcatctttaccatc
tcccagggccagattattgacaactatggaaccaagcctgggaacatcttcctgtgtgtgttccttactcttggagcagccctcactgcattcattaaggcagatctc
cggagacagaaagcaaacaaagaaactcttgagaacaaactccaagaggaggaggaggagagcaacaccagcaaagtgcccactgctgtgtcagaggatca
tctcgattacaaggatgacgacgataagtgagcggccgctcgagtctagagggcccgtttaaacccgctgatcagcctcgactgtgccttctagttgccagccatc
tgttgtttgcccctcccccgtgccttccttgaccctggaaggtgccactcccactgtcctttcctaataaaatgaggaaattgcatcgcattgtctgagtaggtgtcat
tctattctggggggtggggtggggcaggacagcaagggggaggattgggaagacaatagcaggcatgctggggatgcggtgggctctatggcttctgaggcgg
aaagaaccagctggggctctagggggtatccccacgcgccctgtagcggcgcattaagcgcggcgggtgtggtggttacgcgcagcgtgaccgctacacttgcc
agcgccctagcgcccgctcctttcgctttcttcccttcctttctcgccacgttcgccggctttccccgtcaagctctaaatcgggggctccctttagggttccgatttag
tgctttacggcacctcgaccccaaaaaacttgattagggtgatggttcacgtacctagaagttcctattccgaagttcctattctctagaaagtataggaacttcctt
ggccaaaaagcctgaactcaccgcgacgtctgtcgagaagtttctgatcgaaaagttcgacagcgtctccgacctgatgcagctctcggagggcgaagaatctcg
tgctttcagcttcgatgtaggagggcgtggatatgtcctgcgggtaaatagctgcgccgatggtttctacaaagatcgttatgtttatcggcactttgcatcggccgc
gctcccgattccggaagtgcttgacattggggaattcagcgagagcctgacctattgcatctcccgccgtgcacagggtgtcacgttgcaagacctgcctgaaacc
gaactgcccgctgttctgcagccggtcgcggaggccatggatgcgatcgctgcggccgatcttagccagacgagcgggttcggcccattcggaccgcaaggaatc
ggtcaatacactacatggcgtgatttcatatgcgcgattgctgatccccatgtgtatcactggcaaactgtgatggacgacaccgtcagtgcgtccgtcgcgcagg
ctctcgatgagctgatgctttgggccgaggactgccccgaagtccggcacctcgtgcacgcggatttcggctccaacaatgtcctgacggacaatggccgcataa
cagcggtcattgactggagcgaggcgatgttcggggattcccaatacgaggtcgccaacatcttcttctggaggccgtggttggcttgtatggagcagcagacgc
gctacttcgagcggaggcatccggagcttgcaggatcgccgcggctccgggcgtatatgctccgcattggtcttgaccaactctatcagagcttggttgacggcaa
tttcgatgatgcagcttgggcgcagggtcgatgcgacgcaatcgtccgatccggagccgggactgtcgggcgtacacaaatcgcccgcagaagcgcggccgtct
ggaccgatggctgtgtagaagtactcgccgatagtggaaaccgacgccccagcactcgtccgagggcaaaggaatagcacgtactacgagatttcgattccacc
gccgccttctatgaaaggttgggcttcggaatcgttttccgggacgccggctggatgatcctccagcgcggggatctcatgctggagttcttcgcccaccccaactt
gtttattgcagcttataatggttacaaataaagcaatagcatcacaaatttcacaaataaagcatttttttcactgcattctagttgtggtttgtccaaactcatcaat
gtatcttatcatgtctgtataccgtcgacctctagctagagcttggcgtaatcatggtcatagctgtttcctgtgtgaaattgttatccgctcacaattccacacaaca
tacgagccggaagcataaagtgtaaagcctggggtgcctaatgagtgagctaactcacattaattgcgttgcgctcactgcccgctttccagtcgggaaacctgtc
gtgccagctgcattaatgaatcggccaacgcgcggggagaggcggtttgcgtattgggcgctcttccgcttcctcgctcactgactcgctgcgctcggtcgttcggc
tgcggcgagcggtatcagctcactcaaaggcggtaatacggttatccacagaatcaggggataacgcaggaaagaacatgtgagcaaaaggccagcaaaagg
ccaggaaccgtaaaaaggccgcgttgctggcgtttttccataggctccgcccccctgacgagcatcacaaaaatcgacgctcaagtcagaggtggcgaaacccg
acaggactataaagataccaggcgtttccccctggaagctccctcgtgcgctctcctgttccgaccctgccgcttaccggatacctgtccgcctttctcccttcggga
agcgtggcgctttctcatagctcacgctgtaggtatctcagttcggtgtaggtcgttcgctccaagctgggctgtgtgcacgaaccccccgttcagcccgaccgctg
cgccttatccggtaactatcgtcttgagtccaacccggtaagacacgacttatcgccactggcagcagccactggtaacaggattagcagagcgaggtatgtagg
cggtgctacagagttcttgaagtggtggcctaactacggctacactagaagaacagtatttggtatctgcgctctgctgaagccagttaccttcggaaaaagagtt
ggtagctcttgatccggcaaacaaaccaccgctggtagcggtggtttttttgtttgcaagcagcagattacgcgcagaaaaaaaggatctcaagaagatcctttg
atcttttctacggggtctgacgctcagtggaacgaaaactcacgttaagggattttggtcatgagattatcaaaaaggatcttcacctagatccttttaaattaaaa
atgaagttttaaatcaatctaaagtatatatgagtaaacttggtctgacagttaccaatgcttaatcagtgaggcacctatctcagcgatctgtctatttcgttcatcc
atagttgcctgactccccgtcgtgtagataactacgatacgggagggcttaccatctggccccagtgctgcaatgataccgcgagacccacgctcaccggctccag
atttatcagcaataaaccagccagccggaagggccgagcgcagaagtggtcctgcaactttatccgcctccatccagtctattaattgttgccgggaagctagagt
aagtagttcgccagttaatagtttgcgcaacgttgttgccattgctacaggcatcgtggtgtcacgctcgtcgtttggtatggcttcattcagctccggttcccaacg
atcaaggcgagttacatgatcccccatgttgtgcaaaaaagcggttagctccttcggtcctccgatcgttgtcagaagtaagttggccgcagtgttatcactcatgg
ttatggcagcactgcataattctcttactgtcatgccatccgtaagatgcttttctgtgactggtgagtactcaaccaagtcattctgagaatagtgtatgcggcgac
cgagttgctcttgcccggcgtcaatacgggataataccgcgccacatagcagaactttaaaagtgctcatcattggaaaacgttcttcggggcgaaaactctcaa
ggatcttaccgctgttgagatccagttcgatgtaacccactcgtgcacccaactgatcttcagcatcttttactttcaccagcgtttctgggtgagcaaaaacagga
aggcaaaatgccgcaaaaaagggaataagggcgacacggaaatgttgaatactcatactcttcctttttcaatattattgaagcatttatcagggttattgtctcat
gagcggatacatatttgaatgtatttagaaaaataaacaaataggggttccgcgcacatttccccgaaaagtgccacctgacgtc

1

**Supplementary Table 2 – Amino acid sequences of human FLVCR variants used in this study.**

| Variant | Amino acid sequence |
|---|---|
| FLVCR1a (wild-type) | MARPDDEEGAAVAPGHPLAKGYLPLPRGAPVGKESVELQNGPKAGTFPVNGAPRDSLAAASGVLGGPQTPLAPEEETQARLLPAGAGAETPGAESSPLPLTALSPRRFVVLLIFSLYSLVNAFQWIQYSIISNVFEGFYGVTLLHIDWLSMVYMLAYVPLIFPATWLLDTRGLRLTALLGSGLNCLGAWIKCGSVQQHLFWVTMLGQCLCSVAQVFILGLPSRIASVWFGPKEVSTACATAVLGNQLGTAVGFLLPPVLVPNTQNDTNLLACNISTMFYGTSAVATLLFILTAIAFKEKPRYPPSQAQAALQDSPPEEYSYKKSIRNLFKNIPFVLLLITYGIMTGAFYSVSTLLNQMILTYYEGEEVNAGRIGLTLVVAGMVGSILCGLWLDYTKTYKQTTLIVYILSFIGMVIFTFTLDLRYIIIVFVTGGVLGFFMTGYLPLGFEFAVEITYPESEGTSSGLLNASAQIFGILFTLAQGKLTSDYGPKAGNIFLCVWMFIGIILTALIKSDLRRHNINIGITNVDVKAIPADSPTDQEPKTVMLSKQSESAIDYKDDDDK |
| FLVCFR2 (wild-type) | MVNEGPNQEESDDTPVPESALQADPSVSVHPSVSVHPSVSINPSVSVHPSSSAHPSALAQPSGLAHPSSSGPEDLSVIKVSRRRWAVVLVFSCYSMCNSFQWIQYGSINNIFMHFYGVSAFAIDWLSMCYMLTYIPLLLPVAWLLEKFGLRTIALTGSALNCLGAWVKLGSLKPHLFPVTVVGQLICSVAQVFILGMPSRIASVWFGANEVSTACSVAVFGNQLGIAIGFLVPPVLVPNIEDRDELAYHISIMFYIIGGVATLLLILVIIVFKEKPKYPPSRAQSLSYALTSPDASYLGSIARLFKNLNFVLLVITYGLNAGAFYALSTLLNRMVIWHYPGEEVNAGRIGLTIVIAGMLGAVISGIWLDRSKTYKETTLVVYIMTLVGMVVYTFTLNLGHLWVVFITAGTMGFFMTGYLPLGFEFAVELTYPESEGISSGLLNISAQVFGIIFTISQGQIIDNYGTKPGNIFLCVFLTLGAALTAFIKADLRRQKANKETLENKLQEEEEESNTSKVPTAVSEDHLDYKDDDDK |

2

**Supplementary Table 3 – Cryo-EM and model data statistics.**

| | FLVCR1-IF$^{apo}$ (EMD-18334; PDB 8QCS) | FLVCR1-IF$^{choline}$ (EMD-18335; PDB 8QCT) | FLVCR1-IF$^{ethanolamine}$ (EMD-19009; PDB 8R8T) | FLVCR2-IF$^{apo}$ (EMD-18336; PDB 8QCX) | FLVCR2-OF$^{apo}$ (EMD-18337; PDB 8QCY) | FLVCR2-IF$^{choline}$ (EMD-18339; PDB 8QD0) | FLVCR2-OF$^{heme}$ (EMD-18338; PDB 8QCZ) | FLVCR2-IF$^{heme\text{-}choline}$ (EMD-19018) |
|---|---|---|---|---|---|---|---|---|
| **Data collection and processing** | | | | | | | | |
| Magnification | 215,000 | 215,000 | 215,000 | 215,000 | 215,000 | 215,000 | 105,000 | 215,000 |
| Voltage (kV) | 300 | 300 | 300 | 300 | 300 | 300 | 300 | 300 |
| Electron dose ($e^-$/Å$^2$) | 55 | 55 | 55 | 80 | 80 | 57 | 80 | 55 |
| Defocus range (µm) | −1.1 to −2.1 | −1.1 to −2.1 | −1.1 to −2.1 | −1.1 to −2.1 | −1.1 to −2.1 | −1.1 to −2.1 | −1.1 to −2.1 | −1.1 to −2.1 |
| Pixel size (Å) | 0.573 | 0.573 | 0.573 | 0.573 | 0.573 | 0.573 | 0.837 | 0.573 |
| Symmetry imposed | *C*1 | *C*1 | *C*1 | *C*1 | *C*1 | *C*1 | *C*1 | *C*1 |
| Initial particle images (no.) | 3,247,307 | 3,251,081 | 3,086,118 | 3,497,517 | 3,497,517 | 4,790,154 | 5,419,952 | 1,975,710 |
| Final particle images (no.) | 56,664 | 25,873 | 28,295 | 50,167 | 49,532 | 51,511 | 51,266 | 37,044 |
| Map resolution (Å) | 2.9 | 2.6 | 3.1 | 3.1 | 2.9 | 2.8 | 3.1 | 3.1 |
| FSC threshold | 0.143 | 0.143 | 0.143 | 0.143 | 0.143 | 0.143 | 0.143 | 0.143 |
| Map resolution range (Å) | 2.4–6.5 | 2.6–4.9 | 3.0–5.3 | 2.8–5.5 | 2.7–3.8 | 2.6–3.8 | 2.7–5.8 | 2.7–6.0 |
| **Refinement** | | | | | | | | |
| Initial model used | AlphaFold model (AF-Q9Y5Y0-F1) | | | AlphaFold model (AF-Q9UPI3-F1) | | | | |
| Model resolution (Å) | 3.4 | 3.1 | 3.2 | 3.6 | 3.4 | 3.3 | 3.6 | |
| FSC threshold | 0.5 | 0.5 | 0.5 | 0.5 | 0.5 | 0.5 | 0.5 | |
| Model resolution range (Å) | | | | | | | | |
| Map sharpening *B* factor (Å$^2$) | −80 | −30 | −40 | −120 | −100 | −30 | −60 | |
| Model composition | | | | | | | | |
| Non-hydrogen atoms | 3,255 | 3,273 | 3,270 | 3283 | 3,273 | 3,290 | 3,316 | |
| Protein residues | 417 | 418 | 418 | 422 | 421 | 422 | 421 | |
| Ligands | NAG: 1 | NAG: 1<br>CHT: 1 | NAG: 1<br>ETA: 1 | – | – | CHT: 1 | HEM: 1 | |
| Average *B* factors (Å$^2$) | | | | | | | | |
| Protein | 87.45 | 40.28 | 68.08 | 76.93 | 82.55 | 63.63 | 109.15 | |
| Ligand | 144.11 | 68.46 | 93.55 | – | – | 31.72 | 183.4 | |
| R.m.s. deviations | | | | | | | | |
| Bond lengths (Å) | 0.003 | 0.003 | 0.003 | 0.003 | 0.004 | 0.003 | 0.004 | |
| Bond angles (°) | 0.570 | 0.512 | 0.594 | 0.512 | 0.592 | 0.641 | 0.631 | |
| Validation | | | | | | | | |
| MolProbity score | 1.50 | 1.40 | 1.38 | 1.20 | 1.31 | 1.31 | 1.43 | |
| Clashscore | 6.82 | 7.22 | 4.07 | 4.20 | 5.71 | 5.68 | 7.24 | |
| Poor rotamers (%) | 2.5 | 3.1 | 0.6 | 0.6 | 0.0 | 2.8 | 1.4 | |
| Ramachandran plot | | | | | | | | |
| Favored (%) | 97.3 | 98.1 | 96.9 | 99.0 | 98.6 | 99.0 | 98.1 | |
| Allowed (%) | 2.7 | 1.9 | 3.1 | 1.0 | 1.4 | 1.0 | 1.9 | |
| Disallowed (%) | 0.0 | 0.0 | 0.0 | 0.0 | 0.0 | 0.0 | 0.0 | |

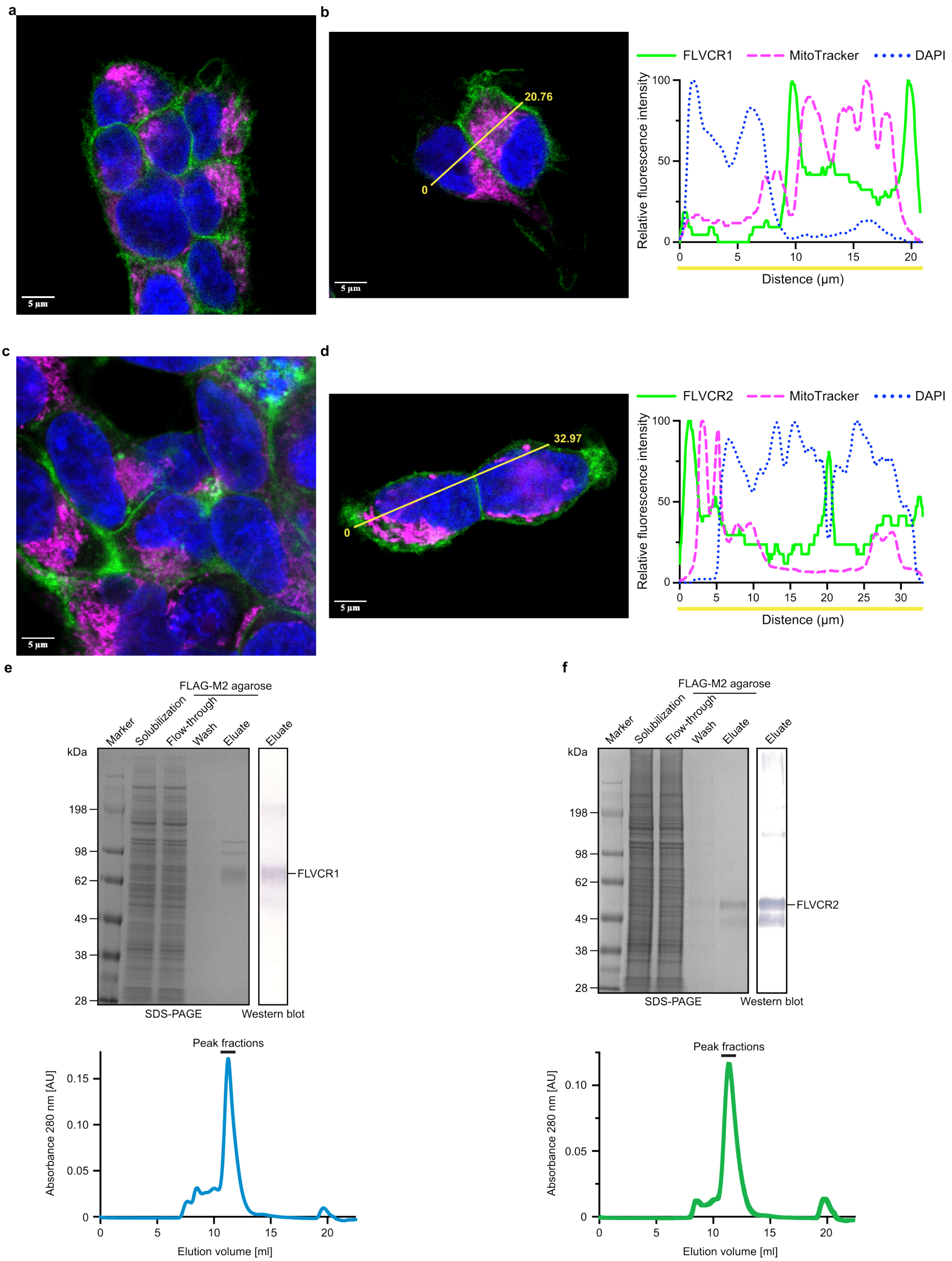


**Supplementary Fig. 1: Purification of human FLVCR1 and FLVCR2.** Representative immunofluorescence images of HEK293 cells overproducing FLAG-tagged wild-type human FLVCR1 (**a**,**b**) and FLVCR2 (**c**,**d**). Both FLVCRs were labelled with ANTI-FLAG® M2-FITC antibody. Mitochondria was labelled with MitoTracker™ Red CMXRos dyes. DNA was stained with DAPI. Distance trace of fluorescence intensities from these dyes within one representative image was analyzed and presented in (**b**) for FLVCR1 and (**d**) for FLVCR2. SDS-PAGE and western blot analysis of FLVCR1 (**e**) and FLVCR2 (**f**) affinity purification (left) and subsequent size-exclusion chromatography (right).

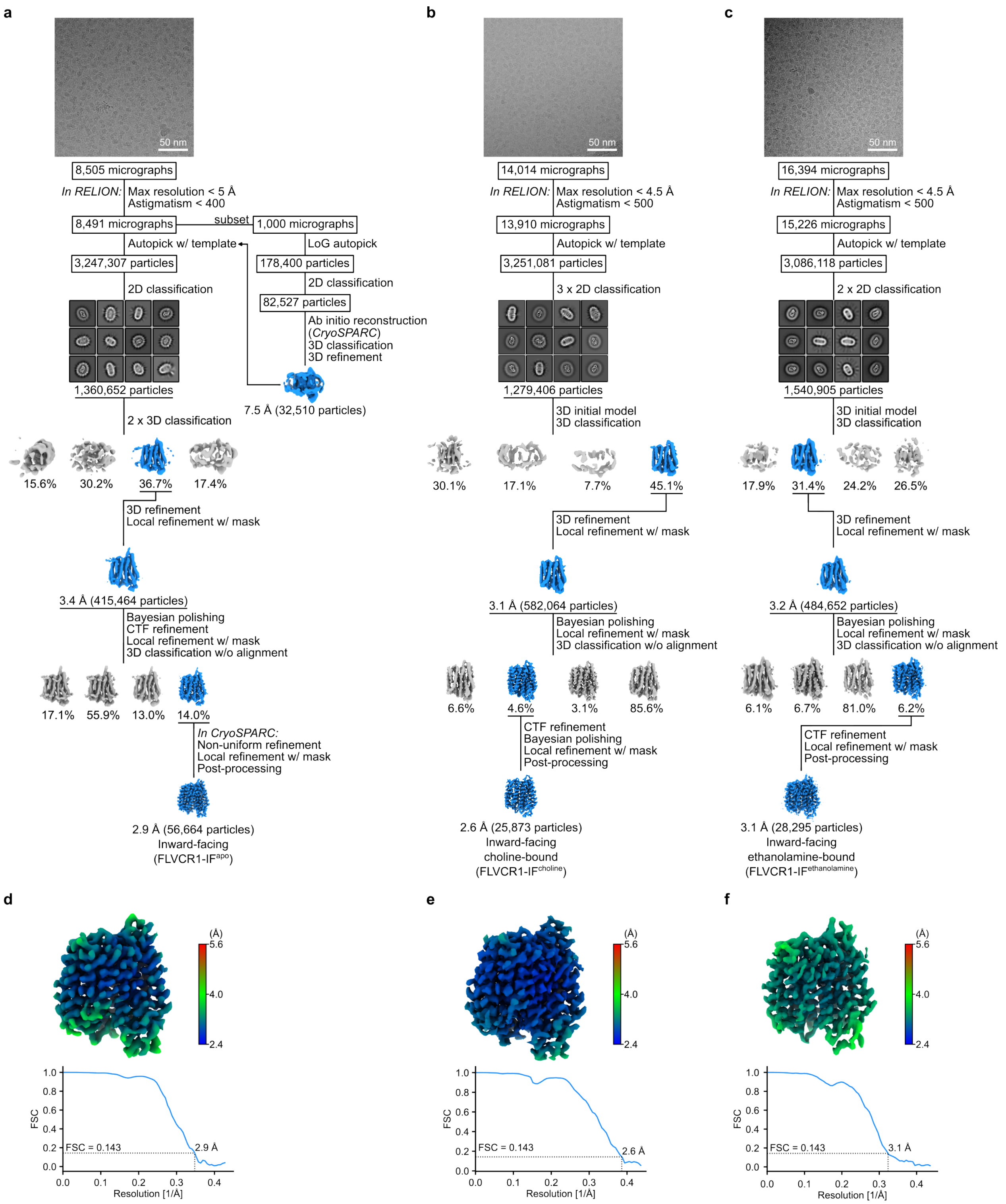


**Supplementary Fig. 2: Single-particle cryo-EM analysis of human FLVCR1.**

**a**-**c**, Summary of the data processing procedure of the as-isolated FLVCR1 sample and FLVCR1 supplemented with choline. **d**-**f**, Local resolution estimation (left) and Fourier shell correlation (FSC) curves (right) of the final cryo-EM maps of FLVCR1-IF$^{apo}$ and -IF$^{choline}$.

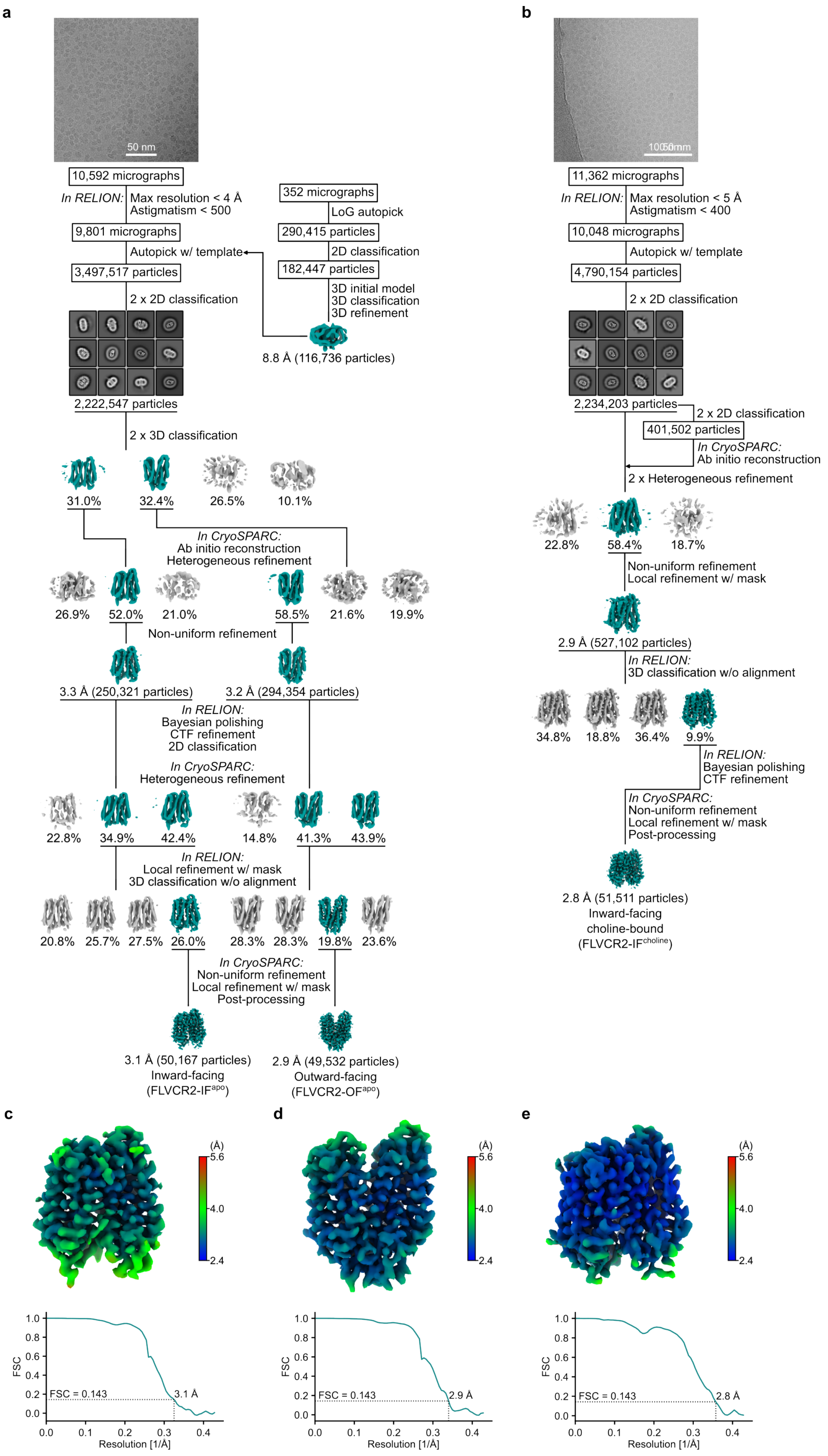
a
50 nm
10,592 micrographs
In RELION: Max resolution < 4 Å
Astigmatism < 500
9,801 micrographs
Autopick w/ template
3,497,517 particles
2 x 2D classification
2,222,547 particles
2 x 3D classification
352 micrographs
LoG autopick
290,415 particles
2D classification
182,447 particles
3D initial model
3D classification
3D refinement
8.8 Å (116,736 particles)
31.0%
32.4%
26.5%
10.1%
In CryoSPARC:
Ab initio reconstruction
Heterogeneous refinement
26.9%
52.0%
21.0%
58.5%
21.6%
19.9%
Non-uniform refinement
3.3 Å (250,321 particles)
3.2 Å (294,354 particles)
In RELION:
Bayesian polishing
CTF refinement
2D classification
In CryoSPARC:
Heterogeneous refinement
22.8%
34.9%
42.4%
14.8%
41.3%
43.9%
In RELION:
Local refinement w/ mask
3D classification w/o alignment
20.8%
25.7%
27.5%
26.0%
28.3%
28.3%
19.8%
23.6%
In CryoSPARC:
Non-uniform refinement
Local refinement w/ mask
Post-processing
3.1 Å (50,167 particles)
Inward-facing
(FLVCR2-IF^apo)
2.9 Å (49,532 particles)
Outward-facing
(FLVCR2-OF^apo)
b
11,362 micrographs
In RELION: Max resolution < 5 Å
Astigmatism < 400
10,048 micrographs
Autopick w/ template
4,790,154 particles
2 x 2D classification
2,234,203 particles
2 x 2D classification
401,502 particles
In CryoSPARC:
Ab initio reconstruction
2 x Heterogeneous refinement
22.8%
58.4%
18.7%
Non-uniform refinement
Local refinement w/ mask
2.9 Å (527,102 particles)
In RELION:
3D classification w/o alignment
34.8%
18.8%
36.4%
9.9%
In RELION:
Bayesian polishing
CTF refinement
In CryoSPARC:
Non-uniform refinement
Local refinement w/ mask
Post-processing
2.8 Å (51,511 particles)
Inward-facing
choline-bound
(FLVCR2-IF^choline)
c
d
e
(Å)
5.6
4.0
2.4
FSC
Resolution [1/Å]
FSC = 0.143
3.1 Å
2.9 Å
2.8 Å

**Supplementary Fig. 3: Single-particle cryo-EM analysis of human FLVCR2.**
**a**,**b**, Summary of the data processing procedure of the as-isolated FLVCR2 and choline-supplemented FLVCR2 samples, respectively. **c**-**e**, Local resolution estimation (top) and FSC curves (bottom) of the final cryo-EM maps of FLVCR2-IF$^{apo}$, FLVCR2-OF$^{apo}$, and FLVCR2-IF$^{choline}$, respectively.

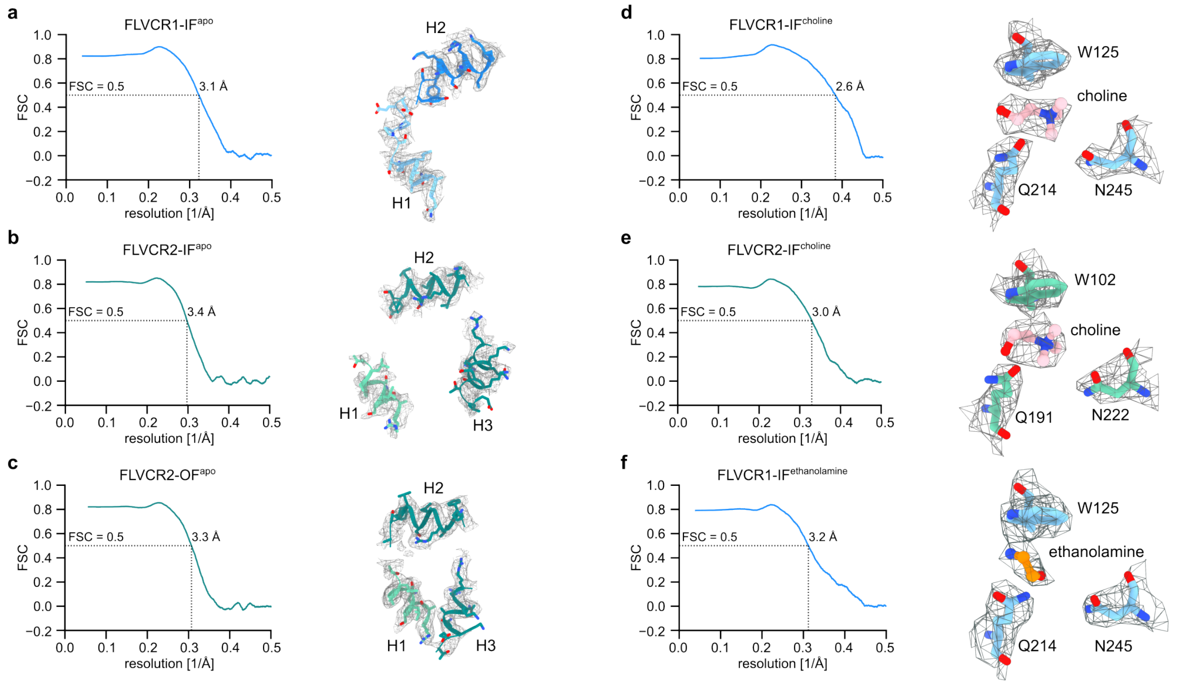


**Supplementary Fig. 4 Cryo-EM density and model fitting.** Cryo-EM map-to-model FSCs of FLVCR1-IF$^{apo}$ (**a**), FLVCR2-IF$^{apo}$ (**b**), FLVCR2-OF$^{apo}$ (**c**),FLVCR1-IF$^{choline}$ (**d**), FLVCR2-IF$^{choline}$ (**e**), and FLVCR1-IF$^{ethanolamine}$ (**f**). The maps for the intracellular helices of the apo states as well as the maps for choline/ethanolamine and the interacting residues of the ligand-bound states are shown.

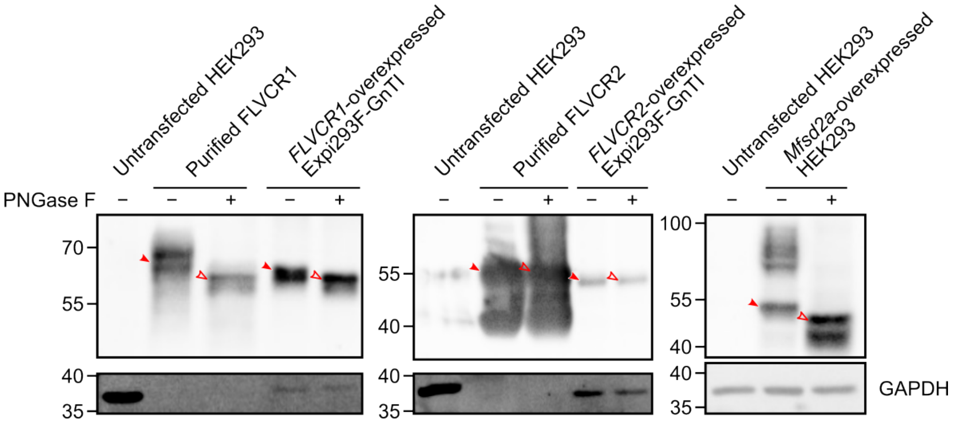


**Supplementary Fig. 5: FLVCR1 deglycosylation assay.**
Western blots of PNGase F treated (+) and untreated (−) samples from cell lysates of Expi293-GnTI cells overexpressing *FLVCR*s or from the purified proteins for cryo-EM analysis. *Mfsd2a*-overexpressed HEK293 cell lysate was used as a control. Filled arrows represent the molecular weight of proteins before deglycosylation and empty arrows represent the molecular weight after deglycosylation.

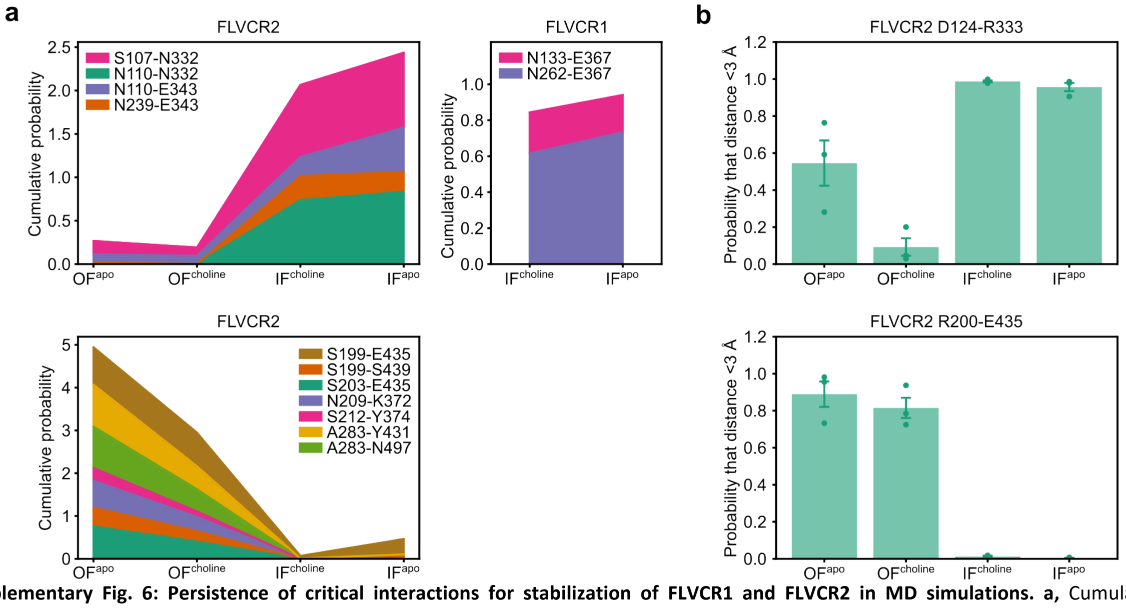


**Supplementary Fig. 6: Persistence of critical interactions for stabilization of FLVCR1 and FLVCR2 in MD simulations. a,** Cumulative probability of inter-domain hydrogen bond formation at the external gate (top) and the internal gate (bottom) with or without choline in the binding cavity. For each pair of residues, the probability was calculated as the fraction of the trajectory in which the distance between the residues remained below 3 Å. **b,** Stability of salt bridges within the external gate (top) and internal gate (bottom) of FLVCR2 in MD simulations. The displayed data represents the mean values obtained from three independent replicas, with error bars representing standard errors of the mean (s.e.m.).

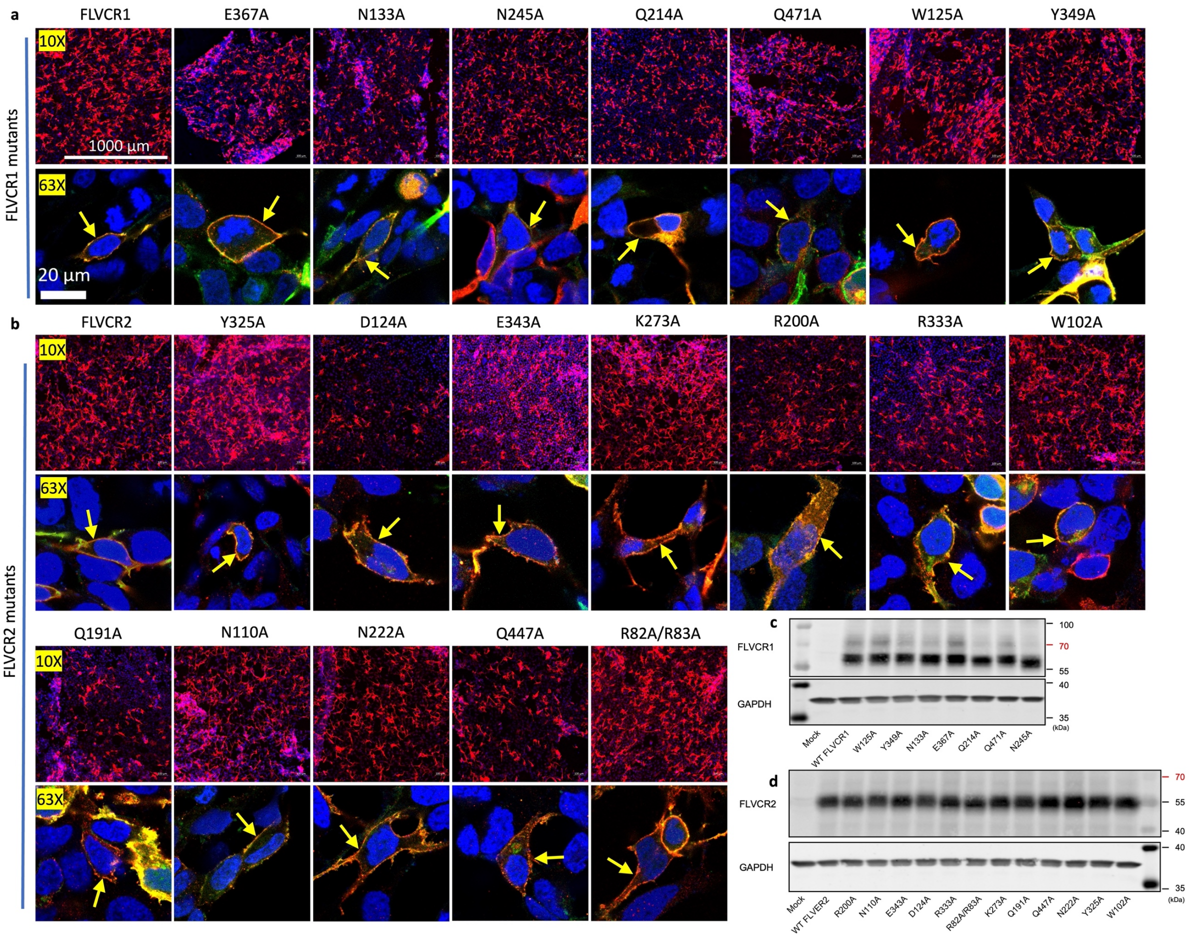


**Supplementary Fig. 7: Immunofluorescent images and western blots of FLVCR mutants.**
Immunofluorescent images of cells overexpressing mutants of *FLVCR1* (**a**) or *FLVCR2* (**b**). Arrows indicate the localization of FLVCRs. Scale bars represent 1000 μm for 10X, and 20 μm for 63X magnification. Western blot analysis of mutated proteins from overexpression in HEK293 cell lysates of FLVCR1 (**c**) or FLVCR2 (**d**).

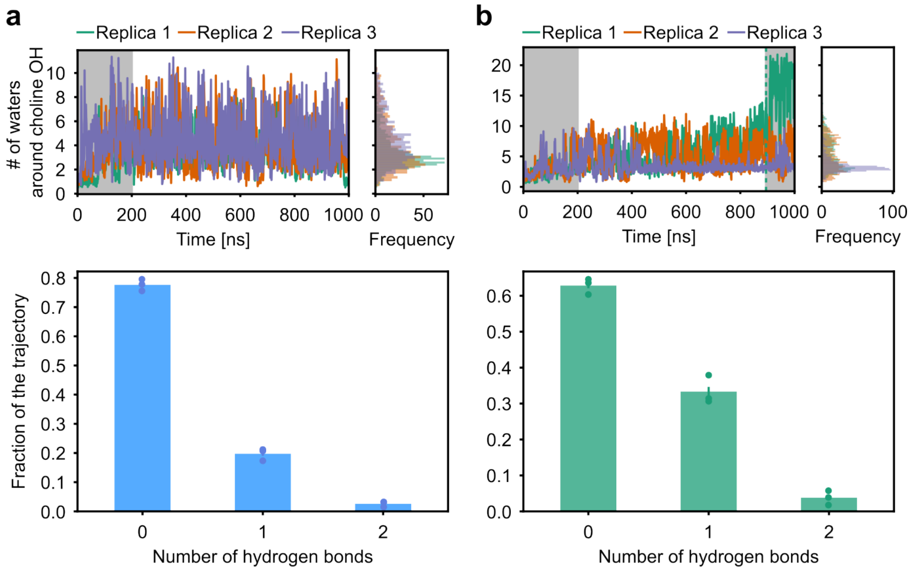


**Supplementary Fig. 8: Hydration of choline in MD simulations.** Interactions between choline and water in the binding site of inward-facing FLVCR1 (**a**) and FLVCR2 (**b**). The upper panels indicate the number of water molecules in the proximity of the hydroxyl group of choline with a cutoff of 5 Å in the $O^{choline}$-$O^{water}$ distance. The first 200 ns of simulations were considered as equilibration and thus not included in the histograms. The last 105 ns of replica 1 in the FLVCR2 simulation were discarded as well due to choline release. The lower panels represent the hydrogen bond formation between waters and the hydroxyl group of choline in the simulations with a distance and angle cut-off of 3 Å and 20 °, respectively. Error bars represent s.e.m.

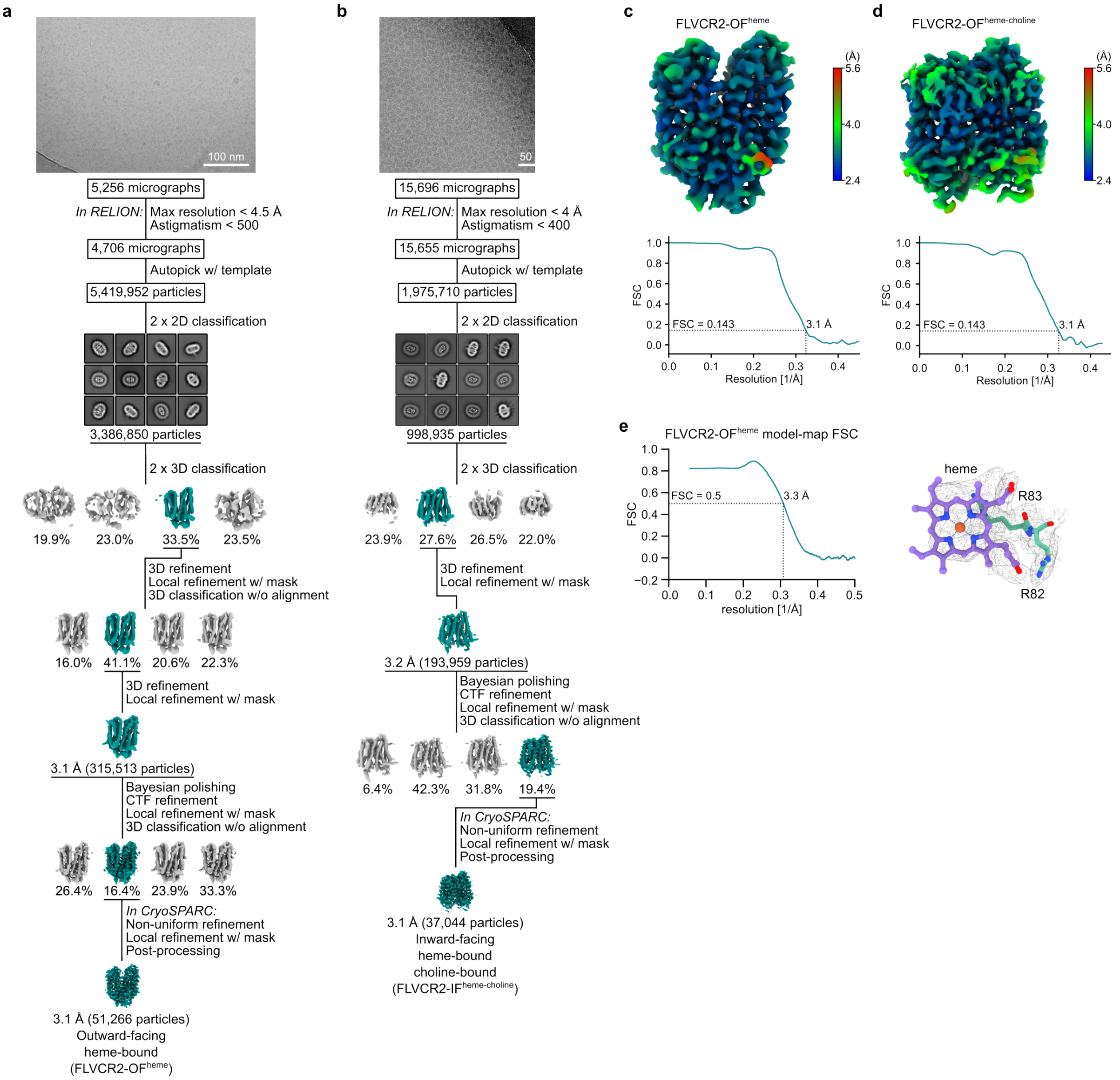


**Supplementary Fig. 9: Single-particle cryo-EM analysis of human FLVCR2 in the presence of heme.**
**a**,**b**, Summary of the data processing procedure of heme-supplemented, and heme-choline-supplemented FLVCR2 samples, respectively. **c**,**d**, Local resolution estimation (top) and FSC curves (bottom) of the final cryo-EM maps of FLVCR2-OF$^{heme}$, and FLVCR2-IF$^{heme-holine}$, respectively. **e**, Cryo-EM map-to-model FSCs of FLVCR2-OF$^{heme}$ and the map fort he bound heme molucule.

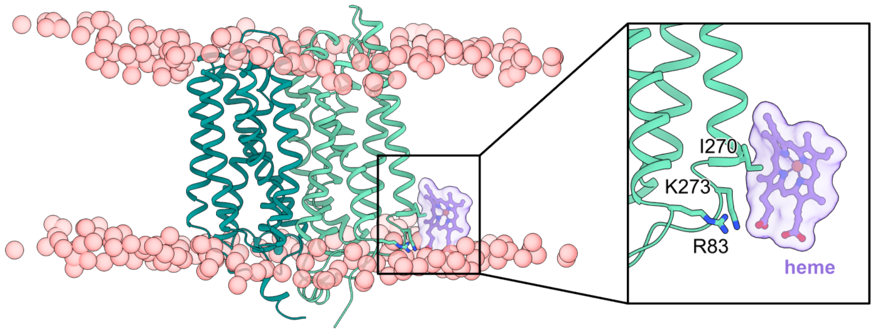


**Supplementary Fig. 10: Snapshot of new, stable interactions formed between heme and FLVCR2 within the lipid bilayer in MD simulations.** The heme molecule is shown in ball-and-stick representation and the phosphates of POPE/POPG are shown as red spheres. The right panel shows the close-up view of the heme molecule and the interacting residues of FLVCR2.

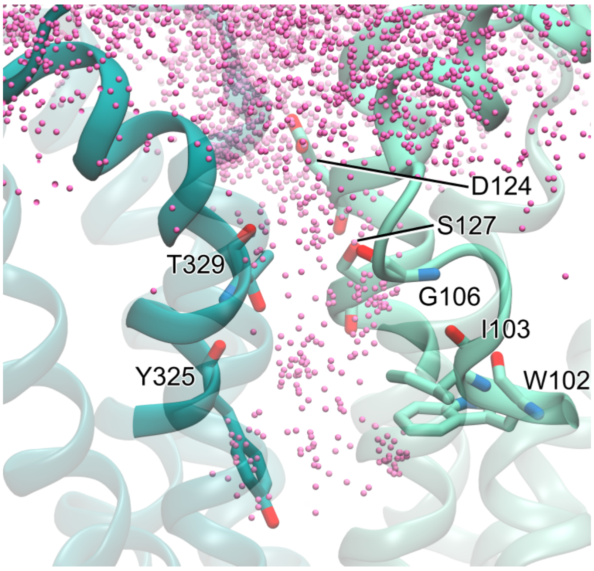


**Supplementary Fig. 11: Choline entry from solution into the outward-facing cavity of FLVCR2 in MD simulations.** Residues that interact with choline frequently are shown as sticks. Pink dots represent choline positions in all frames of one 1-μs trajectory.
